# Absolute abundance unveils *Basidiobolus* as a cross-domain bridge indirectly bolstering gut microbiome homeostasis

**DOI:** 10.1101/2024.12.27.630554

**Authors:** Mitra Ghotbi, Jason E. Stajich, Jason Dallas, Alexander Rurik, Chloe Cummins, Lluvia Vargas-Gastélum, Marjan Ghotbi, Joseph W. Spatafora, Kian Kelly, N. Reed Alexander, Kylie C. Moe, Kimberly C. Syring, Leila Shadmani, Julissa Perez-Marron, Donald M. Walker

## Abstract

The host microbiome is integral to metabolism, immune function, and resilience against pathogens. However, reliance on relative abundance (RA) to estimate host-associated microbiomes introduces compositional biases, while limited tools for absolute abundance (AA) quantification hinder broader applications. To address these challenges, we developed DspikeIn (https://github.com/mghotbi/DspikeIn), an R package paired with a versatile wet-lab methodology for AA quantification. Using RA and AA to compare core microbiome distributions across herpetofauna orders and their natural histories revealed starkly distinct results, driven by aggregate effects, including inherited compositional biases in RA and additional multifactorial influences. Focusing on two closely related *Desmognathus* species demonstrated that AA quantification enhanced resolution in differential abundance analyses and minimized false discovery rates (FDR) when identifying enriched taxa in their gut microbiomes. Keystone taxa identified through network associations also differed between RA and AA data. For example, *Lactococcus* and *Cetobacterium* were core members in Anura and Caudata, while *Basidiobolus* and *Mortierella* were core to Chelonia and Squamata, facilitating host adaptation to diverse environments, insights undetectable with RA data. AA-based network analysis further revealed that removing the *Basidiobolus* subnetwork increased negative interactions, highlighting its role in promoting gut homeostasis through cross-domain connectivity. Despite low redundancy, the *Basidiobolus* node exhibited high betweenness, efficiency, and degree, serving as a critical bridge linking disconnected nodes or modules and indirectly supporting microbiome stability, consistent with Burt’s structural hole theory. DspikeIn represents a transformative tool for microbiome research, enabling the transition from RA to AA quantification and delivering more accurate, consistent, and comparable results across studies.

**Graphical abstract DspikeIn cheatsheet:** 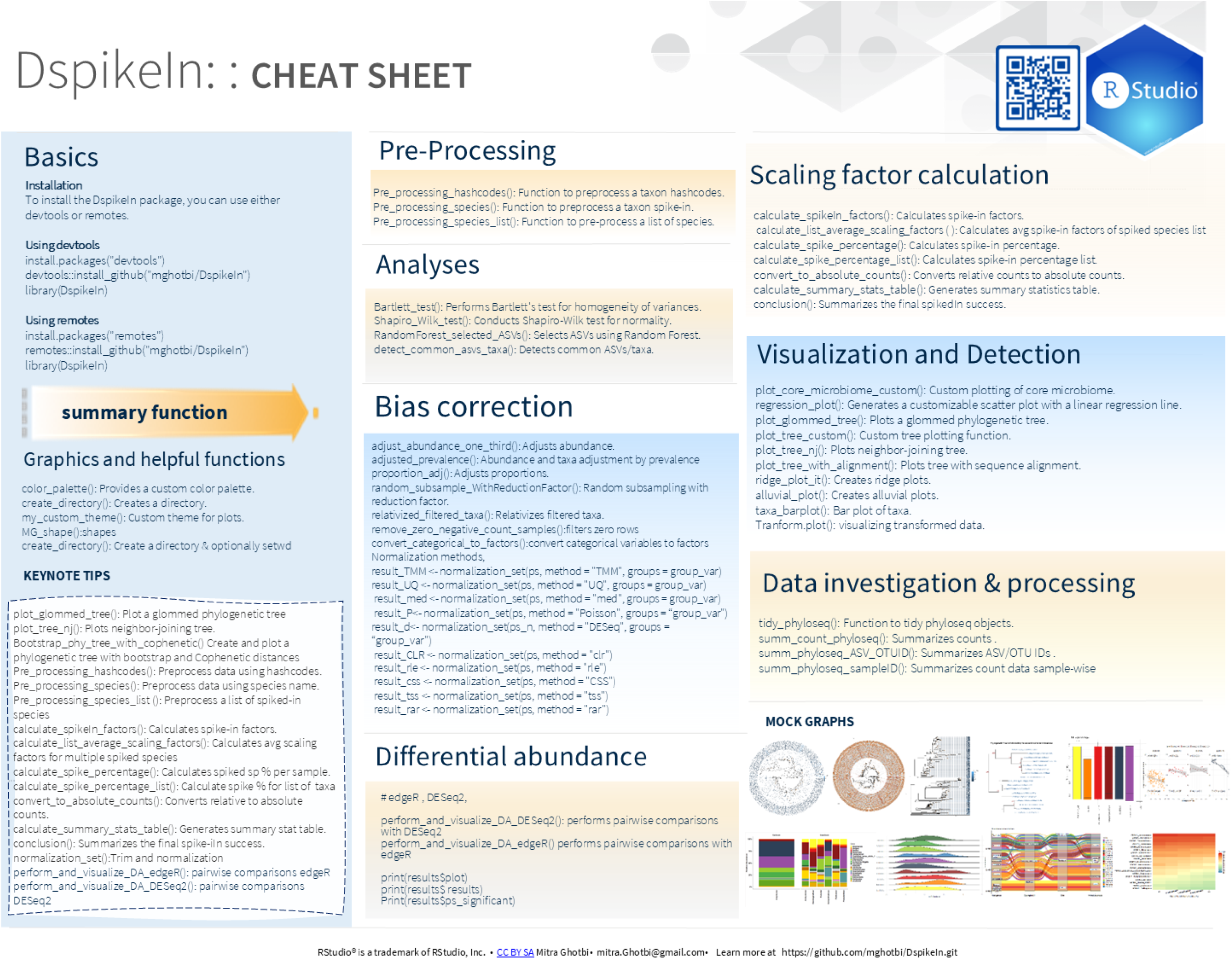

## 1 Introduction

The growing interest in host microbiomes has highlighted a vital role in host metabolism, immune function, and resilience to perturbations [1–3]. Central to these studies is the role of the core microbiome, a stable assemblage of microbial species consistently present across similar hosts or environments. This foundational community offers a reference framework for defining a "healthy" microbiome [4, 5]. Complementing this concept, keystone taxa are species that, despite their relatively low abundance, exert a prominent influence on ecosystem dynamics and functionality [8]. With high connectivity, degree, and betweenness centrality in microbial networks, keystone taxa are crucial for ecosystem stability, and their loss can trigger cascading effects or community reorganization [6, 7]. Among keystone taxa, *Basidiobolus* stands out as a filamentous fungus commonly found in the gastrointestinal tracts of reptiles and amphibians (herpetofauna) [8, 9]. Its unique ability to acquire genes via horizontal gene transfer (HGT) from coexisting gut bacteria [8, 10] further highlights its ecological importance, making it an ideal model for studying bacterial-fungal interactions within herpetofauna gut microbiomes.

Host microbiome studies often employ high-throughput sequencing and multi-omics techniques to unravel complex microbial community dynamics and their interactions with host physiology [11–13]. These approaches have further enhanced the ability to conduct temporal and longitudinal studies, revealing how community shifts influence functional outcomes [14, 15]. However, the data generated through these methods is inherently compositional, reflecting the relative abundance (RA) of microbial communities [16, 17]. RA is constrained to a fixed total of one, meaning an increase in one taxon’s abundance proportionally reduces the apparent abundance of others. This compositional bias undermines the reliability of microbial community studies [18, 19], often introducing spurious negative correlations in association analyses and inflating false-positive rates in differential taxon analyses [19, 20], complicating the interpretation of microbial dynamics and interactions.

To address these limitations, analytical methods, including data transformation techniques [21], and several applications including ALDEx2 [22], ANCOM-BC [18], DEICODE as a QIIME2 plug-in [23], Gneiss [24], Differential Ranking [25], and phylogenetic transform [26], have been developed. These methods employ taxon ratios to address compositional constraints, enhancing the accuracy and relevance of RA datasets [24, 27, 28]. Alongside computational advancements, laboratory techniques such as flow cytometry [29, 30], qPCR [28, 31], digital droplet PCR [28, 32], fluorescence in situ hybridization [33, 34], and fluorescence imaging with correlation spectroscopy [35] have been developed to estimate absolute abundance (AA) from relative counts. Although highly accurate, these techniques are often impractical for large-scale microbiome studies due to scalability limitations [16, 36]. As an alternative, the use of internal standards, or spike-ins, has emerged as a scalable approach for generating AA data from sequencing runs [37–39].

Spike-in approaches involve introducing a known external standard, such as synthetic DNA [36, 38] or non-native microbial whole cells, into a sample [37, 40]. This standard provides a reference point to accurately convert RA into AA values [16]. For whole-cell spike-ins, meticulous selection is essential to ensure that the introduced cells are entirely absent from the studied microbiomes [36, 38]. Whole-cell spike-in methods offer the dual advantage of providing AA and serving as reliable benchmarks to assess variability across key workflow stages, including DNA extraction and amplicon library preparation [16, 38]. Although their adoption is steadily gaining momentum, their application is currently limited to specific ecosystems, such as soil microbiomes [41, 42] and human gut microbiomes [37, 40]. There exists an untapped potential of these methods for broader applications, including the study of gut microbiomes in non-model organisms like herpetofauna. The alarming reality that 40.7% of amphibians and 21.1% of reptiles are threatened with extinction [43, 44] emphasizes the urgency of integrating microbiome research into conservation efforts [45–47]. Investigating herpetofauna microbiomes across diverse natural histories may uncover patterns of dysbiosis and its underlying causes [4, 48, 49], offering valuable insights to inform targeted strategies for mitigating biodiversity loss.

Despite the critical importance of AA for accurately characterizing microbiome dynamics in relation to hosts or environments [17, 50], microbial interactions [40], and microbiome heritability [51], no comprehensive tool has existed to convert RA into AA while addressing potential biases. To address this gap, we developed a wet-lab protocol and the complementary R package DspikeIn for accurate AA quantification. With DspikeIn, we highlight the ecological importance of core and keystone taxa in herpetofauna, showcase how AA enhances the accuracy and relevance of microbiome interpretations, and explore the role of *Basidiobolus* as a core taxon within herpetofauna gut microbiomes. This study addresses three key hypotheses: (a) spiked species retrieval is system-specific rather than following a fixed range; (b) biological interpretations of RA may be biased by compositional constraints; and (c) *Basidiobolus*, as a core member of the herpetofauna gut microbiome, enhances gut microbiome stability through cross-domain mutualistic interactions.

## 2 Methods

### 2.1 Lab techniques and spike volume estimation

Detailed methods are provided in the Supplemental Information (Spike-in Volume Protocol). In brief, *Tetragenococcus halophilus* (bacterial spike; ATCC33315) and *Dekkera bruxellensis* (fungal spike; WLP4642) were selected for spiking gut microbiome samples, as they are absent from wildlife microbiome datasets (GenBank BioProjects: PRJNA1114724, PRJNA1114659). Stock cultures were grown for 72 hours in tryptic soy broth (*T. halophilus*) or potato dextrose broth (*D. bruxellensis*), diluted to OD600 values of 1.0, 0.1, 0.01, and 0.001, and DNA extracted with the Qiagen DNeasy Powersoil Pro Kit. These standards guided spike-in volumes to achieve 0.1–10% of total DNA [69]. To standardize input material, wood frog (*Lithobates sylvaticus*) fecal pellets (3.1 ± 1.6 mg) were processed as 250 µL fecal slurry with and without spiked cells. Nine samples were analyzed to validate the protocol using two metrics: the expected increase in Quantitative PCR (qPCR) cycle threshold (Ct) values proportional to spiked cells and the corresponding increase in copy numbers for *T. halophilus* and *D. bruxellensis*. A synthetic DNA standard curve (16S-V4 rRNA for *T. halophilus* and ITS1 rDNA for *D. bruxellensis*) was used to convert Ct values into log copy numbers for statistical analysis. qPCR was used to compare synthetic DNA of *T. halophilus* and *D. bruxellensis* with DNA extracts and spiked wood frog fecal samples. SYBR Green qPCR assays (20 µL) used Quantabio SYBR Green Fastmix, primers, dsDNAse cleanup, and optimized DNA volumes (*D. bruxellensis*, 1 µL; *T. halophilus*, 3 µL) [52]. Primer pairs (515F/806R for bacteria, ITS1FI2/ITS2 for fungi [8]) were used for amplicon sequencing. Validation involved fecal samples spiked with *T. halophilus* (1874 copies) and *D. bruxellensis* (733 copies), DNA extraction, and Illumina MiSeq sequencing. This streamlined process enabled robust comparisons across sample types [8].

### 2.2 Bioinformatics

Demultiplexed sequencing reads were processed using QIIME2 (version 2023.9) [23]. Paired-end 16S rRNA amplicon reads were assigned to samples based on unique barcodes, with barcode and primer sequences removed using Cutadapt. Reads were merged via FLASH (version 1.2.11) and truncated at 225 bp (forward) and 220 bp (reverse). Two pipelines were used: amplicon sequence variants (ASVs) and operational taxonomic units (OTUs). ASVs were generated using the DADA2 QIIME2 plugin (qiime dada2 denoise-paired), producing unique, error-corrected sequences, while OTUs were clustered de novo at 97% similarity. Taxonomy was assigned using the sklearn classifier with the Silva 138 99% database for the 515F/806R region. Mitochondrial and chloroplast sequences were removed, and a biom file was generated. ITS1 rDNA data underwent a similar process, with ITSxpress [53] used for trimming. Taxonomy was assigned using the UNITE dynamic database (version 10) [54]. This workflow ensured accurate processing, taxonomic classification, and removal of non-target sequences, supporting reliable downstream analyses.

#### 2.2.1 Copy number correction and phylogenetic validation of spiked species

Gene copy number (GCN) correction is not integrated into the DspikeIn package. To account for GCN variability, we utilized the q2-gcn-norm plugin in Qiime2, supported by the rrnDB database (v5.7) [23]. for correcting relative abundance counts. Alternatively, users may apply methods recommended by Louca et al. [55], including PICRUSt, CopyRighter, and PAPRICA. However, due to the inherent variability in rDNA gene copy numbers [56], this correction was not applied to ITS data in this study. Targeted GCN adjustments can still be implemented as needed to prevent overestimating the abundance of specific known fungal taxa.

ASVs from the spiked species were identified using a phylogenetic tree and validated by comparing phylogenetic distances against a reference FASTA file within DspikeIn. These steps, essential for accurate preprocessing, were not required in the OTU approach.

#### 2.2.2 OTU versus ASV approach

To evaluate their suitability for calculating scaling factors to convert RA into AA, OTU and ASV approaches were compared using a spike-in standard. Validation included ten samples: two spiked blanks, two sterile swabs with spiked species, and six randomly selected spiked fecal samples. Both approaches were processed using identical bioinformatics pipelines, and the spiked species retrieval percentage was calculated to assess their accuracy in generating scaling factors.

### 2.3 DspikeIn cheat sheet

The DspikeIn package, built on the phyloseq framework, supports both ASV and OTU approaches, streamlining microbial quantification for diverse experimental setups. It accommodates either a single spike-in taxon or synthetic community taxa with variable or equal spike-in volumes and copy numbers. The package offers a comprehensive suite of tools for AA quantification, addressing challenges through eight core functions: 1) validation of spiked species, 2) data preprocessing, 3) scaling factor extraction, 4) conversion to absolute abundance, 5) bias correction and normalization, 6) system-specific spiked species retrieval, 7) performance assessment, and 8) taxa exploration and filtering. Detailed descriptions of these functions are provided in the supplemental information under "Summary of the DspikeIn Core Features.”

### 2.4 System-specific spiked species retrieval

The system-specific spiked species retrieval threshold was determined by identifying meaningful shifts in key ecological characteristics of the microbial communities. This included assessing the point where spike-in integration (spiked reads per sample) began to affect native microbial metrics such as richness, evenness, beta dispersion, and structure [57]. Together, these metrics established a robust framework for optimal spiked species retrieval.

### 2.5 Relative versus absolute abundance comparisons

Comparative analyses of relative and absolute count datasets were conducted in R (version 4.3.2) using the DspikeIn and vegan packages [58]. Alpha diversity indices (richness and evenness) were calculated from rarefied OTU counts, while beta dispersion was assessed using distance-to-centroid measures to evaluate group variability. Richness, evenness, beta dispersion, and total abundance (total reads) were regressed against spiked species reads using the ‘regression.plot’ function to determine the optimal system-specific spiked species range.

#### 2.5.1 Core microbiome

To assess whether converting to AA reduces spurious discoveries in group comparisons, we analyzed the core microbiome across hosts with varying natural histories (n = 312 samples from wild-caught salamanders, frogs, lizards, turtles, snakes, tortoises, toads, and crocodilians). Using the DspikeIn package and its alluvial_plot function, we visualized the core microbiome across hosts and their natural life histories. To minimize confounding factors in RA vs. AA comparisons, we focused on three closely related host genera (*Desmognathus*, *Eurycea*, and *Plethodon*; n = 120 samples) within three adjacent Tennessee ecoregions. This targeted approach provided a balanced dataset, with results visualized using the taxa_barplot function.

#### 2.5.2 Differential abundance and false discovery rate

Differential abundance results were visualized using volcano plots to compare false discovery rate (FDR) p-values between RA and AA, enabling an evaluation of FDR differences between methods. The Random Forest algorithm (RandomForest_selected_ASVs) identified taxa contributing most to group differences in both datasets, with the top 20 taxa visualized using a ridge plot (ridge_plot_it). For network analyses, differentially abundant taxa were identified to highlight ecologically relevant taxa shaping microbial interactions. Species-specific differences were analyzed using the perform_and_visualize_DA_edgeR function.

#### 2.5.3 Microbial associations

Association networks can lose interpretability when habitat filtering effects become pronounced [59]. To minimize environmental variation and accurately capture taxon-taxon associations, we focused on two closely related salamander species, *Desmognathus imitator* and *D. monticola*, selected for their similar diet, native range, reproductive strategy, and habitat use. This targeted approach reduced false positives and negatives, ensuring a more accurate depiction of microbial interactions. Bacterial and fungal OTUs were rarefied to ∼4,000 reads and merged for network analysis. Cross-domain microbial associations were inferred using the SpiecEasi package (v1.1.2) [60] and visualized in Cytoscape (v3.9.1) [64]. OTUs with >0.05% abundance, present in ≥40% of samples, were included. Network robustness was assessed using StARS with 1,000 subsamples (rep.num = 1000, nlambda = 300), maintaining a lambda of ∼0.49 near the 0.5 stability threshold. Meinshausen-Bühlmann (MB) neighborhood selection was used for network inference, incorporating CLR transformation and sparse inverse covariance estimation to normalize taxonomic variations, reduce biases, and address copy number effects [60]. Additionally, fungal abundances in the cross-domain network were divided by 10 to test shifts in network associations due to higher ITS copy number in fungi compared to 16S gene in bacteria [61].

### 2.6 *Basidiobolus* role and network robustness tests

#### 2.6.1 Phylogenetic diversity and differentially abundant taxa

The phylogenetic diversity of microbial communities associated with *D. imitator* and *D. monticola* was compared to evaluate microbial network stability, using Mean Pairwise Phylogenetic Distance (MPD) as an indicator. Phylogenetic clustering (negative SES) or overdispersion (positive SES) was determined by comparing observed MPD to a null model. The standardized effect size (SES) of MPD was calculated as a Z-score, with the null distribution generated by randomizing taxa labels 999 times using the *picante* package [62]. Phylogenetic diversity between host species was analyzed using one-way ANOVA, followed by pairwise comparisons with Tukey’s HSD test to identify significant group differences.

#### 2.6.2 *Basidiobolus* first connections

The constructed complete network was used to extract nodes and association edges. Communities were identified using the Louvain algorithm for modularity optimization embedded in ‘igraph’ [63]. First, the neighbors of the *Basidiobolus* node were identified using the ‘neighbors’ function. First-order neighbors were directly connected nodes, while second-order neighbors were extracted by iterating through the neighbors of the first-order nodes. Neighboring OTUs were categorized by class, and their distribution was visualized through ‘ggplot2’ [64].

#### 2.6.3 Networks robustness

The modularity network followed conventions described in section 2.5, with nodes of the same color representing membership within the same module. Connectivity patterns classified nodes into four types based on within- and among-module connectivity: keystone taxa (module hubs, highly connected within modules, Zi > 2.5, generalist); network hubs (highly connected across the entire network, Zi > 2.5 and Pi > 0.62, super generalist); connectors (nodes linking different modules, Pi > 0.62, generalist); and peripherals (nodes connected primarily within modules with few external connections, Zi < 2.5 and Pi < 0.62, specialists) [65, 66].

The complete network, constructed using absolute counts, represents the unaltered cross-domain microbial network of the *D. monticola* and *D. imitator* gut microbiomes. Two additional networks were developed: the ‘network and module hubs removed’ (NMHR) network, excluding critical nodes and their edges, and the ‘*Basidiobolus* subnetwork removed’ (BSR) network, where nodes and edges associated with the *Basidiobolus* genus were removed. Negative edge weights, which hindered the calculation of network topological properties, were excluded. A one-way ANOVA using the ggpubr package [67] was performed to compare the networks’ topological properties, as detailed in Table S1.

To evaluate network robustness, the fractional size of the largest connected component (LCC) was regressed against the fraction of removed nodes [68, 69]. Constraint values measured the extent to which a node’s connections were concentrated among immediate neighbors, with high values indicating redundancy and low values reflecting ‘structural holes’ where nodes act as bridges between otherwise disconnected groups [70]. Redundancy for each node was calculated as the sum of edges among its neighbors, excluding direct links to the node [70, 71]. Effective size, estimating unique connections, was calculated as node degree minus redundancy, while efficiency was determined by dividing effective size by degree to measure the proportion of unique connections. Efficiency, redundancy, and betweenness centrality were regressed against node degree using scatter plots [85], and nodes in the top 40% for redundancy or efficiency were identified and labeled for their contributions to network structure and stability. Metrics were calculated using igraph [72] and custom scripts, with visualizations created in ggplot2 [64].

## 3 Results

### 3.1 Phylogenetic validation and rationale for the selection of OTUs over ASVs

For the ASV approach, phylogenetic distances between spiked species ASVs and the reference Sanger sequence were validated prior to preprocessing (Fig. S1A). The OTU-based approach was selected for its 100% retrieval efficiency of spiked species using De Novo Closed-Reference Clustering at 97% similarity in control samples. Additionally, the OTU method yielded a higher proportion of samples within the desired spiked species retrieval range (Fig. S1B-C).

### 3.2 Range of spiked species retrieval

To determine the acceptable range of spiked species retrieval, various biological metrics were plotted and regressed against spiked species reads. In the bacterial dataset, notable shifts occurred when samples exceeded 20% for the distance to centroid metric and 30% for evenness. Beyond 20% spiked species reads, distance to centroid decoupled from spiked species reads, while evenness showed an increasing correlation, indicating that retrieval above 30% is unreliable for estimating scaling factors and absolute abundance (Fig. 1A-B). In the ITS rDNA dataset, similar trends were observed, though evenness remained stable up to 40% spiked species retrieval, providing greater flexibility (Fig. S2A-D).

**Figure 1.**
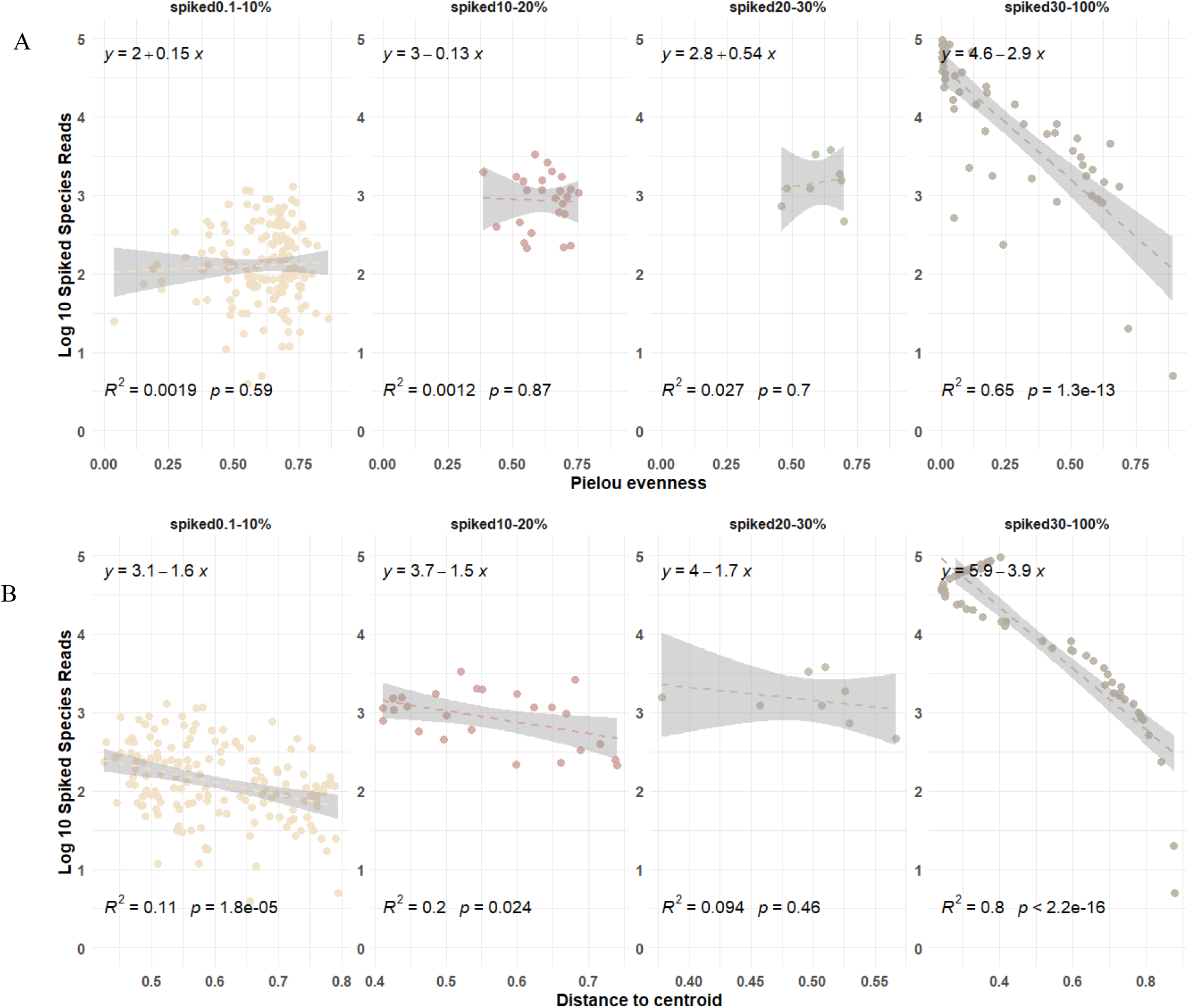
Linear relationship between the log10-transformed reads of bacterial spiked species (*Tetragenococcus halophilus*) and two community metrics, Pielou’s evenness (A) and distance to centroid (B) across varying ranges of spiked species retrieval percentages. Each panel shows a different retrieval range: 0-10%, 10-20%, 20-30%, and 30-100%. Regression lines, *R*^2^ values, and *p* values indicate the strength and significance of associations within each range (Fungal pair results are displayed in Figure S2).

### 3.3 Relative versus absolute abundance comparisons

#### 3.3.1 Core microbiome dynamics

The AA alluvial plot revealed distinct core microbiome distribution patterns linked to host orders and natural history traits, such as diet, habitat, and reproduction. In contrast, the RA core microbiome and mycobiome distributions, exhibited different patterns. Genera like *Lactococcus*, *Cetobacterium*, and *Parabacteroides* (bacteria), along with *Basidiobolus* and *Mortierella* (fungi), were more abundant across herptile traits in the AA plot, whereas RA plots showed genera more evenly distributed across traits and ecoregions (Fig. 2A-B, Fig. S3A-B). *Lactococcus* and *Cetobacterium* dominated aquatic and semi-aquatic hosts, such as anurans and chelonians, while adapting to terrestrial squamates. AA effectively captured herpetofauna natural histories: *Lactococcus* was linked to carnivorous diets, while *Cetobacterium* was associated with insectivory.

**Figure 2.**
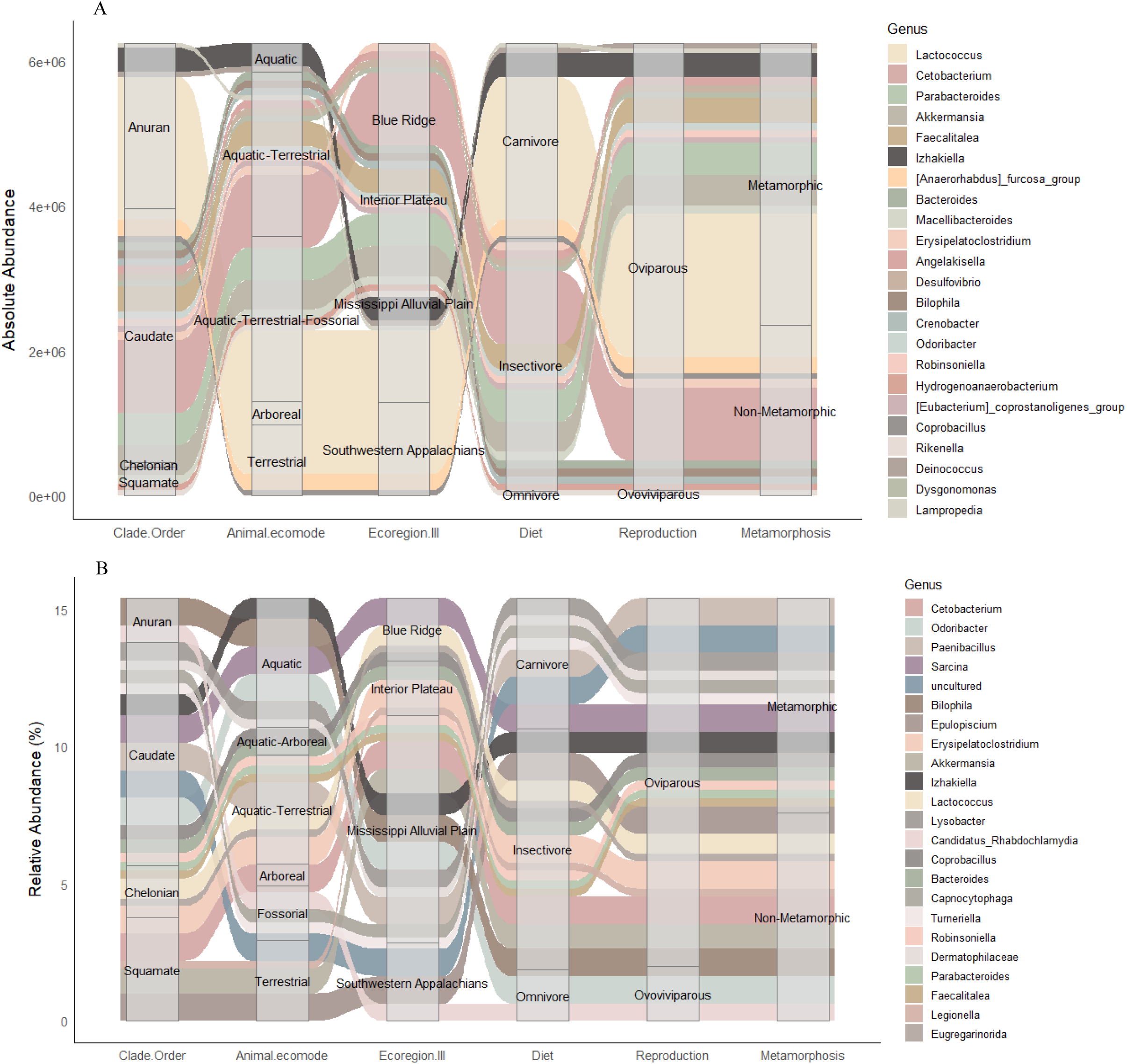
Core microbiome distribution at genus level across herpetofauna orders, shown by absolute abundance (A) and relative abundance (B) in alluvial plots. Plots illustrate the flow of shared bacteria across host natural history, including host order, ecomode, ecoregion, diet, reproduction type, and metamorphosis status. Similar color-code was used for genus in both plots for easy traceability.

To account for aggregate effects, the dataset was subsetted to three closely related host genera (*Desmognathus*, *Eurycea*, *Plethodon*) within three adjacent ecoregions, enabling accurate AA vs. RA comparisons. Bacterial and fungal community compositions differed notably between AA and RA across hosts and ecoregions (Fig. 3A-D). The gut microbiome of Blue Ridge *Desmognathus* was dominated by diverse taxa such as *Biolophila*, *Akkermansia*, and *Parabacteroides*, while *Lactococcus* and *Biolophila* were primary contributors in Interior Plateau *Eurycea*. RA-based profiling showed consistent patterns, with *Odoribacter* and *Robinsoniella* more prevalent in both *Eurycea* and *Plethodon*. Using AA, *Basidiobolus* emerged as the most abundant fungal genus across hosts, except for Eurycea in the Interior Plateau and Southwestern Appalachians, a pattern undetected with RA (Fig. 3B-D).

**Figure 3.**
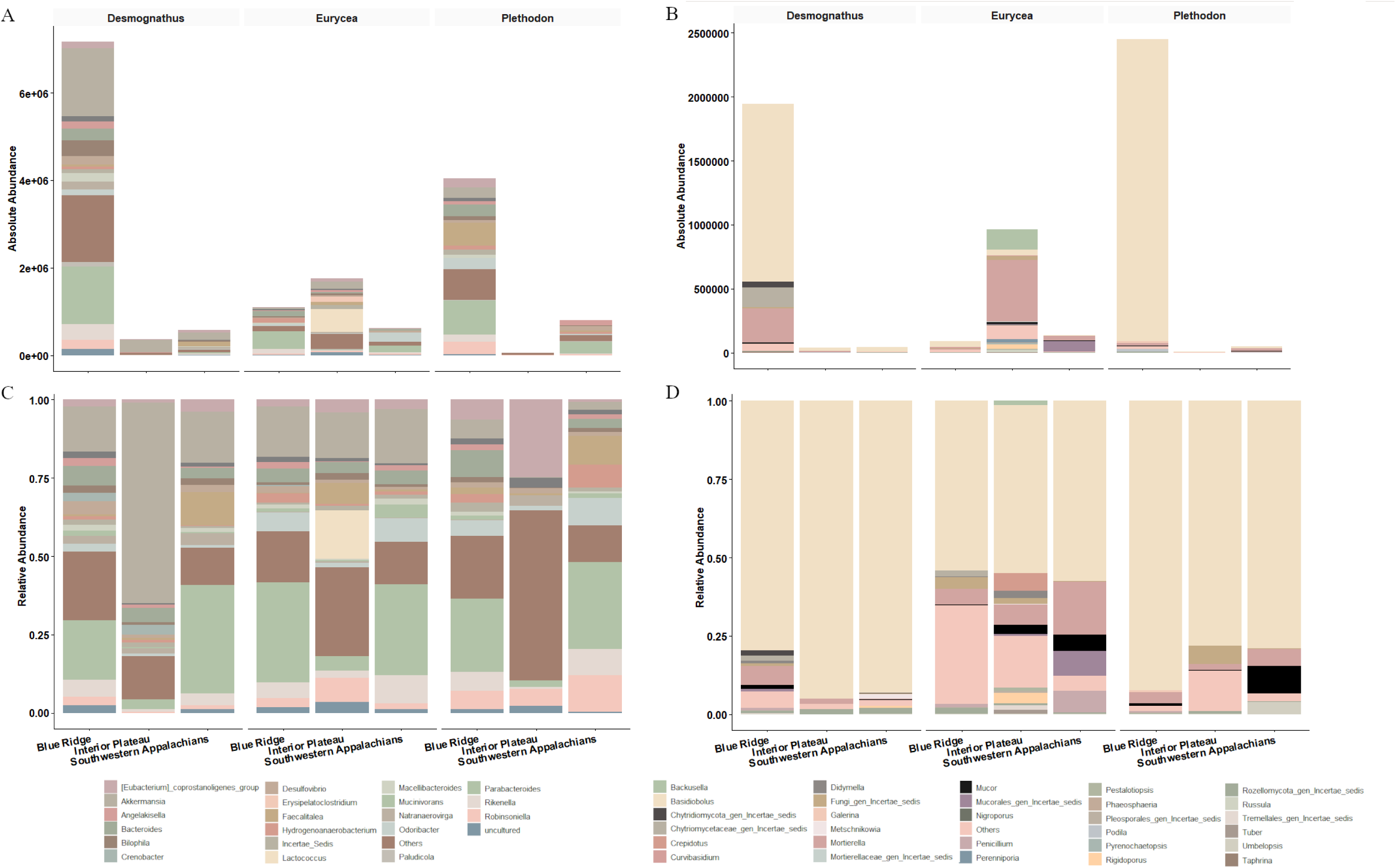
Variation in dominant gut bacteria (A and C) and fungi (B and C) among three genera (*Desmognathus*, *Eurycea*, and *Plethodon*) in the Plethodontidae family across three adjacent ecoregions (Blue Ridge, Interior Plateau, and Southwestern Appalachians) in Tennessee, USA. Absolute abundance is shown in A and B while relative abundance is shown in C and D.

Random Forest classification identified host-associated bacterial and fungal OTUs, highlighting distinct taxonomic variations among the *Desmognathus*, *Plethodon*, and *Eurycea* genera when comparing RA and AA datasets (Fig. S4A-B, Fig. S5A-B).

#### 3.3.2 Differential abundance and False Discovery Rate (FDR)

Volcano plots comparing differential abundance between two closely related *Desmognathus* species with similar natural histories showed that AA provided greater accuracy than RA by eliminating the FDR (Fig. S6A-B, S7A-B). Comparisons of bacterial and fungal differential abundance further highlighted the limitations of RA in reflecting microbial abundance relative to underlying factors (Fig. S6C-D, S7C-D). Differential abundance analyses revealed that Firmicutes genera were dominant in *D. monticola* based on AA but appeared less prominent in RA. For the fungal community, AA showed that Ascomycota and Basidiomycota genera were more prominent in *D. monticola* compared to *D. imitator*.

#### 3.3.3 Microbial cross-domain associations

Comparisons of microbial networks constructed using RA and AA highlighted the limitations of RA in accurately capturing taxon-taxon associations. Key differences were observed in topological properties, including network shape and metrics such as node count (AA: 308, RA: 151), degree (AA: 7.43, RA: 4.09), edge betweenness (AA: 354.02, RA: 210.31), and within- and between-module connectivity, whereas, module count (AA: 18, RA: 19), modularity (AA: 7.43, RA: 7.11), and weight (AA: 0.13, RA: 0.11) were similar (Fig. 4A-E). The network and module hubs identified also differed between the RA- and AA-based networks. Despite rarefying reads and using the Spiec-Easi package to mitigate GCN biases, we further adjusted fungal abundances by dividing them by 10 to account for fungal rDNA GCN being about an order higher than bacterial GCN and the fungal spike-in *Dekkara*. This adjustment had minimal impact on network topology (Fig. S8A-C).

**Figure 4.**
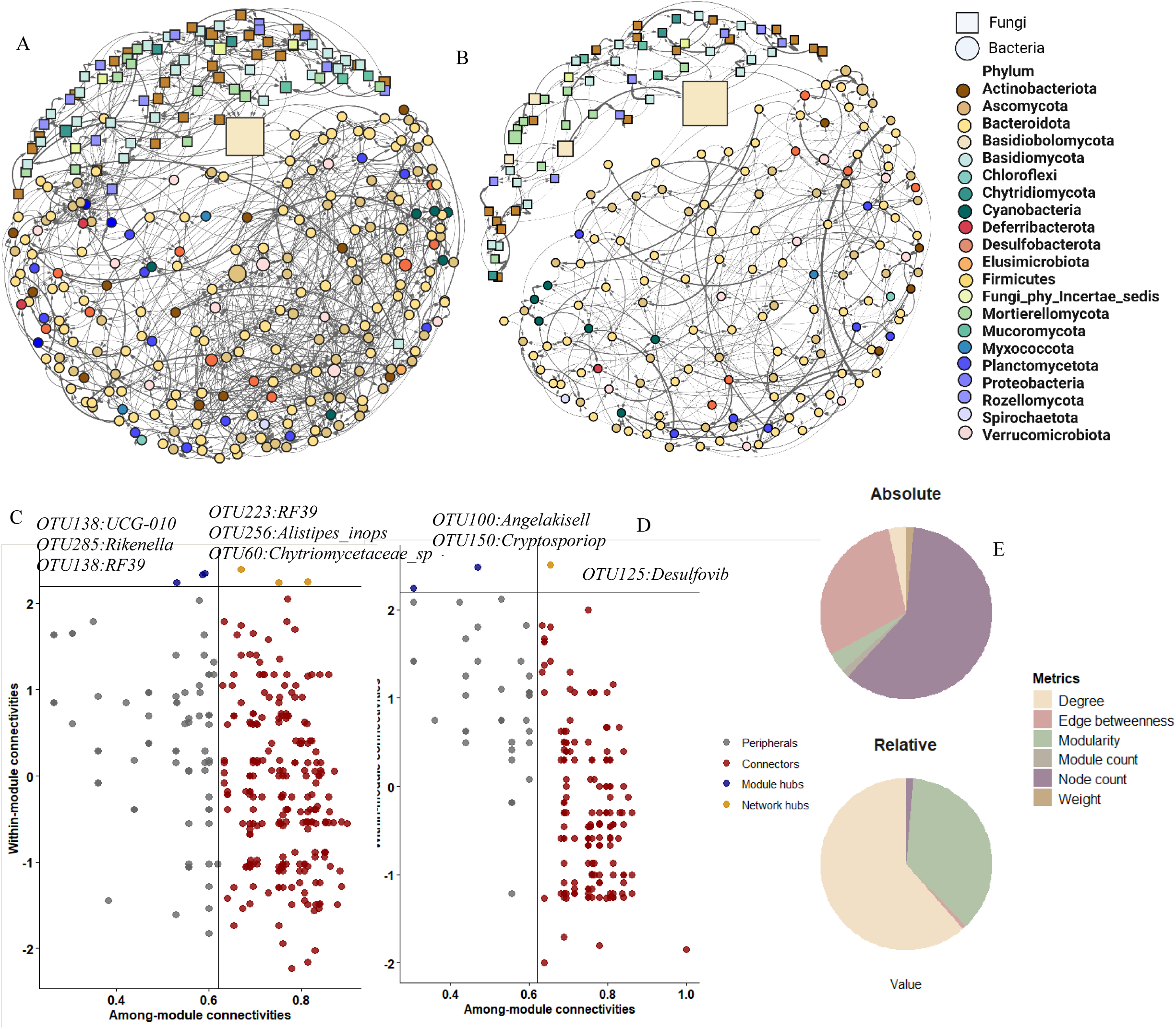
Cross-domain network associations and connectivity profiles of gut microbial communities from two salamander species (*Desmognathus monticola* and *D. imitator*) from the same ecoregion were constructed based on absolute (A) versus relative (B) abundance. Network nodes were categorized based on their connectivity in the network, including within-module and among-module connections, for absolute (C) and relative (D) counts. The nodes are color-coded to represent OTUs associated with specific phyla in the association network. Node size corresponds to the relativized abundance of each phylum. Node shapes indicate domains: circles represent Bacteria, and squares represent Fungi. Edge colors depict the weight of microbial associations, with gray indicating positive associations. Mean topographic properties of relative count-based versus absolute count-based networks are shown (E). Terminology can be found in Table S1.

### 3.4 *Basidiobolus* role and network robustness

#### 3.4.1 Phylogenetic structure and differential abundance

Phylogenetic structure analysis revealed diverse fungal and bacterial communities in both *D. imitator* and *D. monticola*, reflecting stable microbial associations [73] (Fig. S6E-F, S7E-F). Positive SES values indicated phylogenetic overdispersion (competition), while negative values reflected clustering (environmental filtering) [74]. Environmental filtering was significant for fungi in three samples, whereas most bacterial samples exhibited overdispersion. Mean Pairwise Distance (MPD) analysis showed similar fungal (p = 0.30) and bacterial (p = 0.16) phylogenetic diversity between *D. monticola* and *D. imitator* (Fig. S6G-H, S7G-H).

#### 3.4.2 *Basidiobolus* neighbors in network analysis

The complete network comprised 18 modules /communities (Fig. 5A, S9D). In the gut microbiome network, Bacteroides, Bacilli, Clostridia, and Verrucomicrobiae were identified as primary bacterial neighbors of the *Basidiobolus* node. Secondary bacterial neighbors included Actinobacteria, Bacteroides, Clostridia, Bacilli, Desulfovibrionia, Gammaproteobacteria, Vampirivibrionia, and Verrucomicrobiae (Fig. S9E).

**Figure 5.**
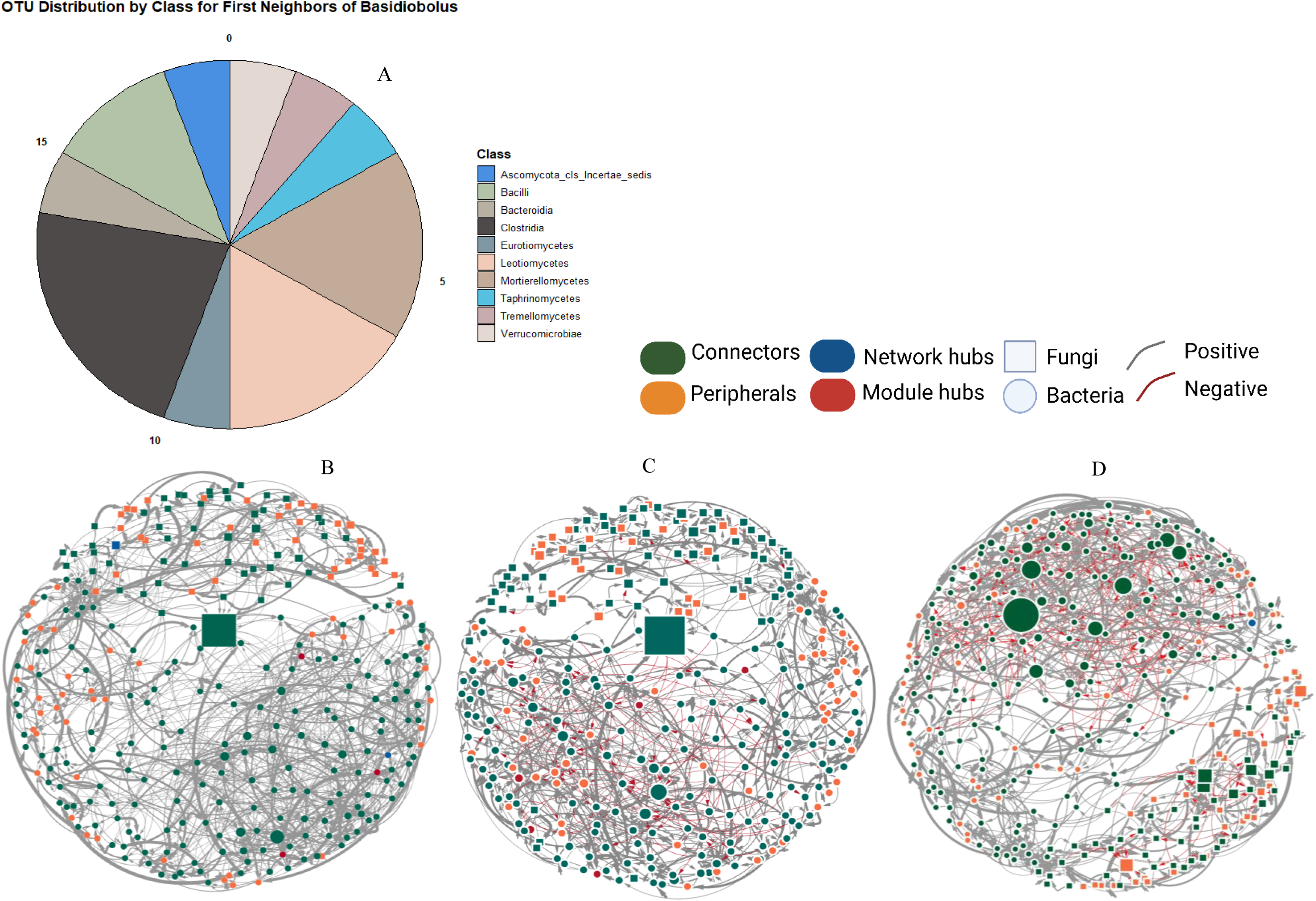
The distribution of OTUs by class for *Basidiobolus*’ first neighbors in gut microbial networks of *D. monticola* and *D. imitator* are shown in (A), with pie chart segments representing microbial classes and segment size reflecting OTU proportions. Cross-domain network associations of *D. monticola* and *D. imitator* gut microbiomes using absolute counts without modification (complete) (B), after removing network and module hubs (C), after removing the *Basidiobolus* subnetwork (D) are shown. Networks are color-coded by connectivity patterns (see Fig. S8 for modularity networks). Similar colors indicate taxa with comparable roles, node shapes differentiate bacteria (circles) from fungi (squares), and edge colors and thickness represent association weights, with gray indicating positive associations (Terminology can be found in Table S1).

#### 3.4.3 Network topology and stability test

To investigate the ecological role of *Basidiobolus* in the herptile gut microbiome, we analyzed three absolute AA-based networks for *D. monticola* and *D. imitator*: the complete network, NMHR, and BSR networks. The removal of the *Basidiobolus* subnetwork (BSR) had a greater impact on network shape and diameter compared to the removal of network and module hubs (NMHR), which also altered modularity and connectivity patterns (Fig. 5B-D, S9A-C, S10A-C). Metrics such as local efficiency, harmonic centrality, and closeness were similarly affected in both BSR and NMHR networks (complete metrics analyses in Fig. S11) (Fig. 6A-E). The complete network displayed exclusively positive interactions, while NMHR showed an 11% increase in negative interactions and BSR resulted in a 14.4% increase. Connectivity and modularity networks emphasized the role of individual taxa in maintaining network integrity, with *Basidiobolus* identified as a connector taxon facilitating cross-domain inter-module interactions (Fig. 5B-D, S9A-C).

**Figure 6.**
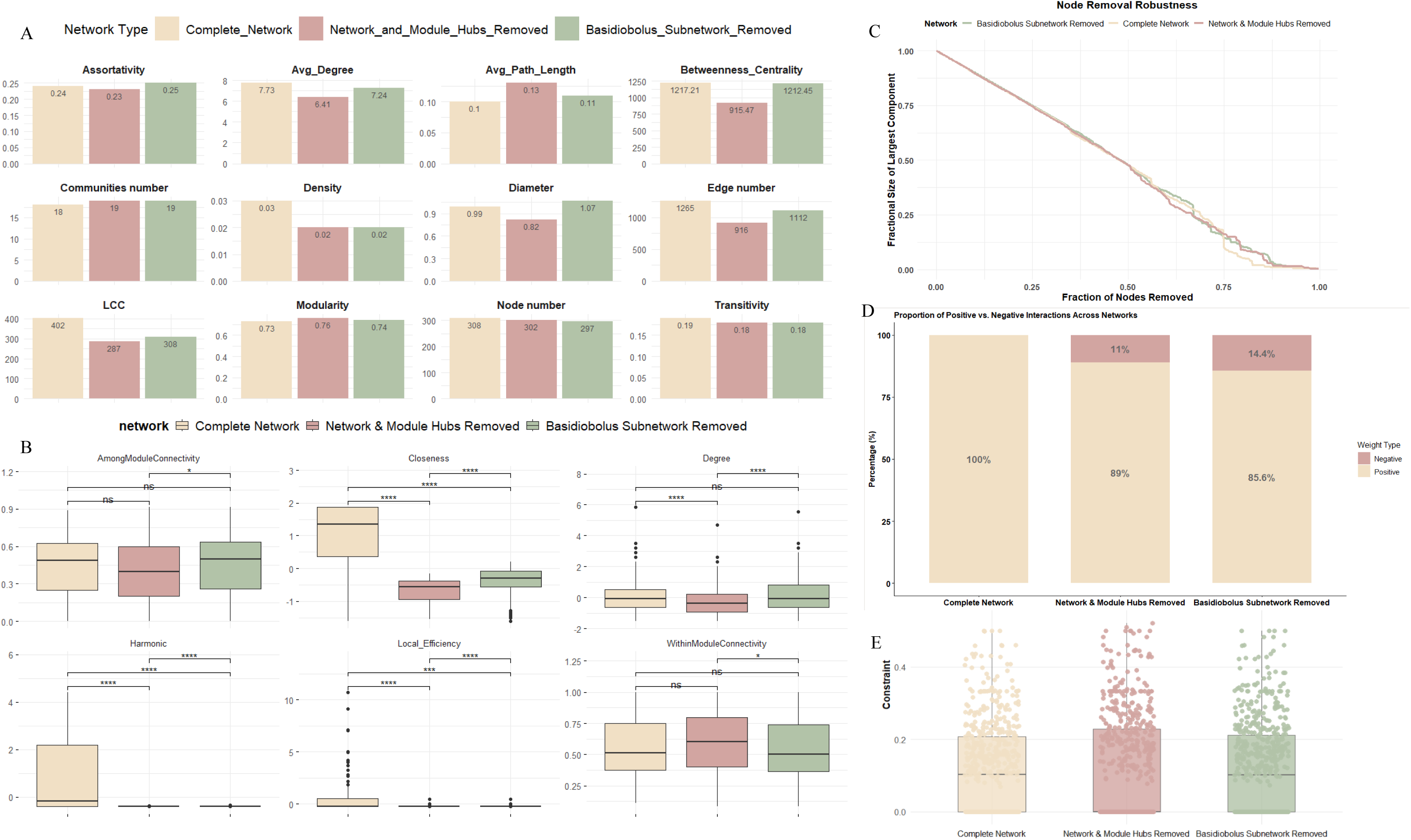
Comparison of average network topology metrics from the three cross-domain networks (refer to Figure 5) (A). Node-based topology was analyzed using ANOVA (B), with significant pairwise differences indicated by asterisks (ns = non-significant; significance codes: p < 0.05 ‘\**’, p < 0.001 ‘\*\*\**’). Network robustness was evaluated by examining the effects of randomly removing nodes (C). Proportions of positive versus negative interactions across network types are shown (D), along with average constraint values (E). Network types are represented as follows: Complete Network (cream), Network with Module Hubs Removed (dark pink), and *Basidiobolus* Subnetwork Removed (light green). Terminology is provided in Table S1.

The fractional size of the LCC decreased smoothly with the fraction of excluded nodes in both NMHR and BSR scenarios (Fig. 6C). Key robustness metrics, including LCC and modularity, showed minimal changes in BSR, with a moderate shift in average path length. High redundancy highlighted the network’s resilience by providing functional buffering through overlapping connections. In the complete network, species exhibited varying redundancy levels (mean: 1.2, median: 1.0). Nodes like OTU256 and OTU268 occupied key positions in the top-right quadrant, serving as highly connected and locally stable hubs. Connectivity metrics, such as betweenness and degree, revealed that *Basidiobolus* (OTU69) exhibited both high betweenness and degree, underscoring its role as a central hub and bridge within the network (Fig. 7A-C). Positioned in the low redundancy, high betweenness, high degree, and high efficiency region, *Basidiobolus* (OTU69) played a less critical role in network stability but served as a key bridge between domains and communities.

**Figure 7.**
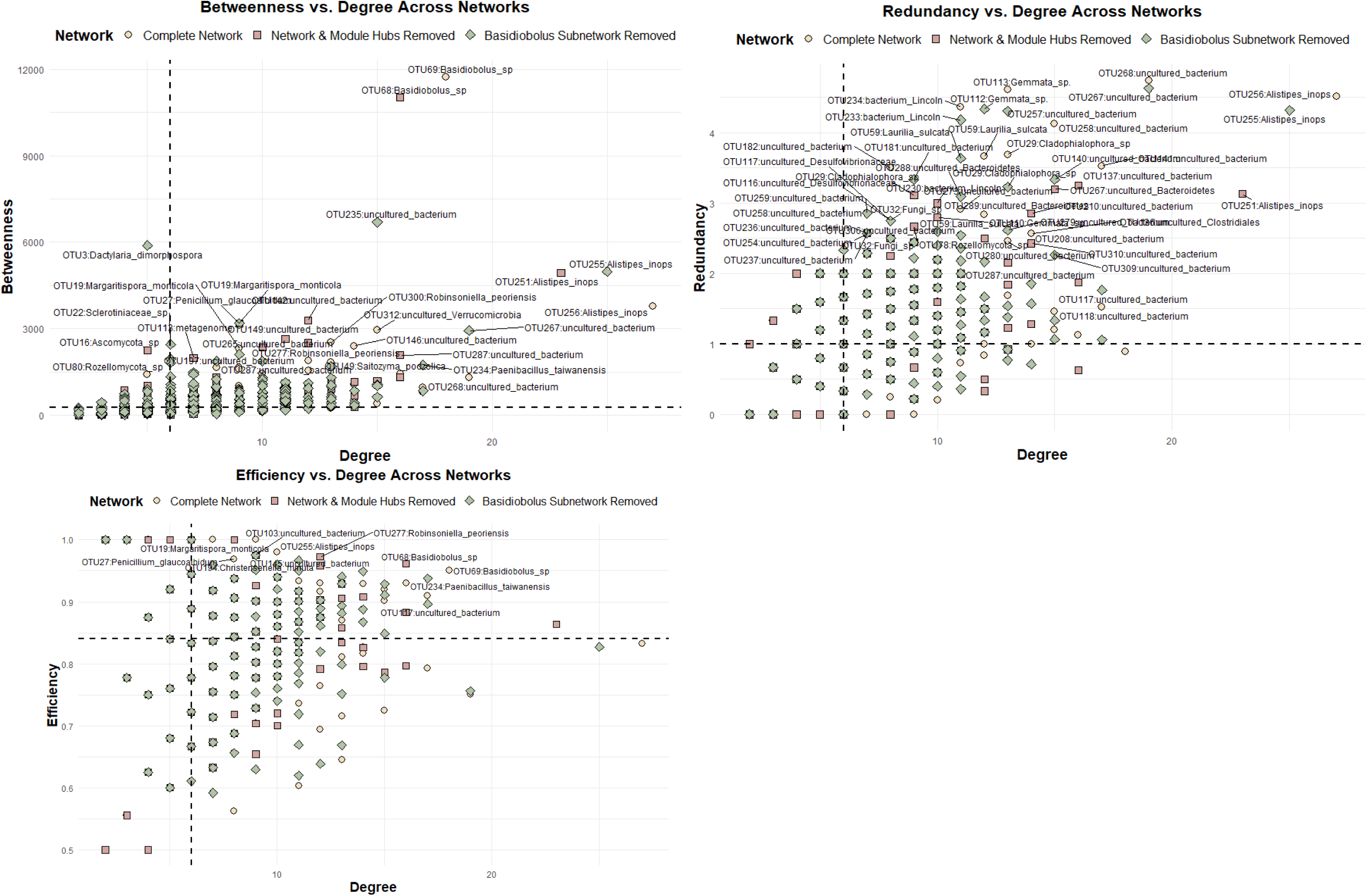
Network stabilities of the gut microbiomes from *Desmognathus imitator* and *D. monticola*. Stability metrics include node-level betweenness plotted against degree (A), node-level redundancy (B), and efficiency (C) plotted against degree, with dividing lines representing the medians of each axis. Redundancy was calculated as the sum of edges among a node’s neighbors, and efficiency as the ratio of effective size to degree. Nodes in the top 40% for redundancy, efficiency or betweenness were labeled on figures for their roles in network structure and stability. Terminology is provided in Table S1

## 4 Discussion

### 4.1 DspikeIn for converting to absolute and addressing biases

DspikeIn is a consolidated R package paired with a versatile wet-lab methodology, validated in a non-model animal gut microbiome system. Sequencing workflows are prone to biases, which can emerge at any stage, from PCR primer selection to DNA extraction, amplification, and sequencing [75–77]. Incorporating whole-cell spike-ins provides a robust benchmark for assessing the entire workflow, from sample storage and DNA extraction to library preparation and the refinement of computational thresholds and analyses [28, 36]. DspikeIn facilitates the exploration of host microbiomes by incorporating whole-cell spike-ins and AA quantification, uncovering patterns and interactions often misrepresented or absent in traditional RA analyses.

The debate surrounding the use of OTUs versus ASVs remains ongoing, with contrasting perspectives highlighting their respective advantages and limitations [78, 79]. Using whole-cell spike-ins, OTUs outperformed ASVs in converting RA to AA, achieving nearly 100% recovery of spiked species in positive controls, making OTUs the preferred approach. Intraspecific and occasional intragenomic variability in the ITS region further supports clustering sequences for species-level resolution [80]. Therefore, we opted for clustering approaches over ASVs and did not apply copy number correction for ITS data. For users employing ASVs, we recommend first evaluating mean phylogenetic distances and/or comparing ASV phylogenetic relationships to the positive control (e.g., a Sanger FASTA file). ASVs from spiked species should then be merged for scaling factor computation, a step easily performed using DspikeIn.

DspikeIn models the relationship between microbial metrics and spiked species reads, expanding the previously established 0.1–10% cutoff range [40]. This study demonstrated acceptable spiked species retrieval thresholds of 30% for bacteria and 40% for fungi. These findings emphasize that optimal spiked species retrieval ranges are system-specific and should be tailored to the unique characteristics and metrics of each study. Metrics such as evenness, richness, and distance to centroid are sensitive to perturbations [81]. While detecting a threshold for the fungal community, we applied a 30% threshold in downstream analyses to ensure greater biological relevance.

### 4.2 Relative versus absolute abundance comparisons

Examining microbial communities across populations is essential for identifying stable core microbiota components and distinguishing them from transient species [82]. In this study, we systematically compared the accuracy and efficiency of AA and RA in interpreting biological outcomes. Our findings demonstrated that AA significantly reduced spurious discoveries, enhancing the precision of identifying core microbiome and keystone taxa in relation to the natural history of diverse herpetofauna. Network analyses further revealed notable differences in topology and keystone taxa between AA and RA networks. The core microbiome forms a foundational community that underscores critical contributors to essential ecological roles and functions [5, 82]. Similarly, keystone taxa play a pivotal role in sustaining community structure and integrity [83, 84]. These keystone species, whether acting individually or within guilds, profoundly influence microbiome architecture and functionality, transcending their RA or spatial and temporal distributions [85].

AA revealed specific genera that were masked or overrepresented in RA data, providing clearer insights into microbial associations with host traits and natural history. This improvement aligns with findings across diverse hosts and habitats, including quantitative analyses and host-associated systems [17, 29, 36, 37]. By eliminating compositional biases inherent in RA-based methods, AA has consistently enhanced microbial community profiling and interaction analysis [30, 40]. This refinement enables more accurate assessments of microbial dynamics, including their associations with host phenotypes and environmental variables, and advances our understanding of ecological and pathological relationships [28, 29, 40]. Using AA, we identified *Lactococcus* and *Cetobacterium* as consistently dominant genera across herpetofauna. These findings suggest that *Lactococcus* and *Cetobacterium* are core microbiome members with broad ecological adaptability. Their true abundance and associations with host natural history, often obscured in RA data, highlight the importance of AA for accurate biological interpretation.

The prevalence of *Lactococcus* and *Cetobacterium* in the gut microbiomes of amphibians may suggest functionalities similar to those observed in Giant Amazonian Fish *(Arapaima gigas)* [86] and zebrafish (*Danio rerio*) [87]. In *A. gigas*, *Cetobacterium* constituted 55–87% of the gut microbiome, playing key roles in host immune regulation and enhancing microbial community resilience, particularly against spring viremia of carp virus [86]. Similarly, in zebrafish, *Cetobacterium* demonstrated the capacity to synthesize vitamin B12, which stabilized the gut microbiome and significantly improved host resistance to *Aeromonas hydrophila* infection [87]. Likewise, *Lactococcus* species contribute to gut microbiome diversity and exert probiotic effects, as demonstrated in swine [88]. Accurately revealing core microbiome distributions using AA across host natural histories improves our understanding of key taxa and their roles in promoting host health and resilience.

### 4.3 *Basidiobolus* role in gut microbiome association networks

We identified *Basidiobolus* as a dominant member of the core microbiome and a highly connected keystone taxon within the gut microbiome networks of herpetofauna clades. This filamentous fungus, commonly found in the gastrointestinal tracts of a wide range of herpetofauna [89, 90], plays a pivotal role in maintaining the stability and functionality of herpetofauna gut ecosystems [8]. With its presumed acquisition of bacterial genes through horizontal gene transfer, akin to other anaerobic gut fungi [91], *Basidiobolus* demonstrates remarkable adaptability within the gut environment [10]. This unique capacity underscores its significance as a model organism for studying bacterial-fungal interactions in herpetofauna gut microbiomes.

In the gut microbiome network, the *Basidiobolus* node demonstrated direct positive associations with Bacteroides, Bacilli, Clostridia, and Verrucomicrobiae, suggesting potential mutualistic relationships with these coexisting microbial taxa. Collectively, these taxa play key roles in gut health by facilitating digestion, maintaining microbial balance, and enhancing mucosal and metabolic functions [92, 93].

Recent research highlights the critical role of mycobiota in microbiota equilibrium and gut health [94] with microbial adaptability such as siderophore production, being central to these interactions. Siderophores play a critical role in bacterial iron acquisition under iron-limited conditions, promoting bacterial growth, shaping microbial community dynamics, and regulating virulence in pathogenic bacteria [95, 96]. Compounds like schizokinen and catecholates, produced by bacteria such as *Bacillus* spp., further promote microbial balance and host iron absorption [97–99], underscoring their critical contribution to gut health and nutrition.

Bacteroides engages in both symbiotic and competitive interactions within the gut microbiome [100] and exhibiting antigrowth and antivirulence activities against fungi such as *Candida albicans* [101]. A hallmark of *Bacteroides* (phylum Bacteroidota) is its ability to degrade diverse polysaccharides through polysaccharide utilization loci, which encode enzymes that capture, transport, and break down polysaccharides, producing metabolites like short-chain fatty acids essential for gut health and homeostasis [92, 102]. Similarly, *Clostridium* species play a vital role in gut homeostasis, supporting gut function by producing beneficial metabolites like butyrate [93, 103]. These species are indispensable for modulating physiological, metabolic, and immune processes, interacting with resident microbes, and supporting gut health throughout the host’s lifespan [103, 104].

The essential role of *Basidiobolus* in mediating mutualistic interactions within the gut microbiomes of *D. monticola* and *D. imitator* was evident in the loss of positive cohesion following the removal of the *Basidiobolus* subnetwork. This removal increased negative interactions to 14.4%, compared to 11.0% after removing network or module hubs. Understanding network interfaces is crucial for interpreting biotic factors driving secondary metabolite production, which is vital for maintaining host homeostasis and overall health [105, 106]. From a biological perspective, microbial community interactions can be classified as positive or negative, each driven by distinct mechanisms. Positive interactions arise from processes like cross-feeding, co-aggregation in biofilms, co-colonization, niche overlap, or cooperative strategies that promote microbial synergy. Conversely, negative interactions often stem from dynamics like amensalism, prey-predator relationships, competition, or other antagonistic mechanisms that disrupt microbial coexistence and community balance [65, 107–109]. In microbial networks, these interactions shape stability. A proposed framework [6] suggests that positive interactions can enhance stability or, under certain conditions, lead to destabilization. The loss of a single species can cascade through the network, threatening stability [6, 110]. This aligns with the stress-gradient hypothesis, where positive interactions provide temporary resilience under stress but increase fragility if critical connections are lost [6, 110]. A diverse community resists disturbances more effectively, as competition among well-adapted species limits overgrowth, making diversity and redundancy crucial for microbial resilience [105, 111]. Although *Basidiobolus* exhibited low redundancy (limited connection overlap) and did not directly affect global stability metrics, its high efficiency, betweenness, and degree underscored its role as a cross-domain bridge, enhancing localized interaction efficiency between bacteria and fungi.

Mechanisms that reduce interaction intensity enhance microbial association stability, with redundancy replacing a few strong interactions with numerous weaker ones [109, 111]. The observed increase in negative interactions following the removal of the *Basidiobolus* subnetwork highlighted the network’s heightened vulnerability to its connections. Despite low redundancy and constraints, the *Basidiobolus* node exhibited high connectivity, reinforcing its role as a bridge linking otherwise unconnected nodes or modules, consistent with Burt’s structural hole theory [70]. While *Basidiobolus* may not have directly influenced gut microbiome balance and homeostasis, it likely supported these processes indirectly through its bridging role.

## 5. Conclusion

DspikeIn marks a major advancement in microbiome research by enabling the transition from RA to AA quantification. This shift enhances benchmarking, producing results that are more consistent, comparable, and reliable across studies. AA effectively minimized FDR, improved accuracy, and strengthened links between microbial dynamics and the host’s natural life history. The removal of *Basidiobolus* was linked to increased negative interactions in the gut microbiome networks of two closely related salamanders, reinforcing its likely role in preserving cross-domain mutualistic interactions. By leveraging AA, we further identified *Basidiobolus* as a crucial bridge within the network, characterized by its significant betweenness, degree, and efficiency in facilitating these interactions. This may be attributed to horizontal gene transfer from first-neighbor bacteria, such as Bacilli. These species are known for secreting siderophores, and/or mutualistic associations with Bacteroides, Clostridia, and Verrucomicrobiae, which may involve nutrient cross-feeding, facilitate digestion, and maintain microbial balance. These results highlight the versatility of the DspikeIn tool and underscore the necessity of AA quantification, paving the way for identifying key biological interactions in future research.

## Data availability

Data and DspikeIn package deposited on GitHub at https://github.com/mghotbi/DspikeIn. The raw sequence data are deposited in NCBI Sequence Read Archive under BioProject number PRJNA1202922.

## Acknowledgment

We extend our sincere appreciation to Dr. Kaitlyn Murphy (Department of Biology, Middle Tennessee State University) for meaningful discussions on the manuscript.

