## Supplemental for "Absolute abundance unveils *Basidiobolus* as a cross-domain bridge indirectly bolstering gut microbiome homeostasis"

Supplemental Information

Spike-in volume Protocol,

The species Tetragenococcus halophilus (bacterial spike; ATCC33315) and Dekkera bruxellensis (fungal spike; WLP4642-White Labs) were selected as taxa to spike into gut microbiome samples as they were not found in an extensive collection of wildlife skin (GenBank BioProjects: PRJNA1114724, PRJNA 1114659) or gut microbiome samples (PRJNA PRJNA1202922). Stock cell suspensions of both microbes were grown in either static tryptic soy broth (T. halophilus) or potato dextrose broth (D. bruxellensis) for 72 hours then serially diluted and optical density (OD600) determined on a ClarioStar plate reader. Cell suspensions with an optical density of 1.0, 0.1, 0.01, 0.001 were DNA extracted using the Qiagen DNeasy Powersoil Pro Kit. These DNA isolations were used as standards to determine the proper spike in volume of cells to represent 0.1-10% of a sample (Rao et al., 2021b) Fecal pellets (3.1 ± 1.6 mg; range = 1 – 5.1 mg) from an ongoing live animal study using wood frogs (Lithobates sylvaticus) were used to standardize the input material for the development of this protocol. A total of (n=9) samples were used to validate the spike in protocol. Each fecal sample was homogenized in 1mL of sterile molecular grade water then 250uL of fecal slurry was DNA extracted as above with and without spiked cells.

Two approaches were used to evaluate the target spike-in of 0.1-10%, the range of effective spike-in percentage described in (Rao et al., 2021b), including 1) an expected increase of qPCR cycle threshold (Ct) value that is proportional to the amount of spiked cells and 2) the expected increase in copy number of T. halophilus and D. bruxellensis in spiked vs. unspiked samples. A standard curve was generated using a synthetic fragment of DNA for the 16S-V4 rRNA and ITS1 rDNA regions of T. halophilus and D. bruxellensis, respectively. The standard curve was used to convert Ct values into log copy number for statistical analyses (detailed approach in(Romer et al., 2022;

Walker et al., 2019)) using the formula y = -0.2426x + 10.584 for T. halophilus and y = -0.3071x + 10.349 for D. bruxellensis, where x is the average Ct for each unknown sample.

Quantitative PCR (qPCR) was used to compare known copy numbers from synthetic DNA sequences of T. halophilus and D. bruxellensis to DNA extractions of T. halophilus and D. bruxellensis independently, and wood frog fecal samples with and without spiked cells. SYBR qPCR assays were run at 20ul total volume including 10ul 2X Quantabio PerfeCTa SYBR Green Fastmix, 1ul of 10uM forward and reverse primers, 1ul of ArcticEnzymes dsDNAse master mix clean up kit, and either 1ul of DNA for D. bruxellensis or 3ul for T. halophilus. Different volumes of DNA were chosen for amplification of bacteria and fungi due to previous optimization of library preparation and sequencing steps (Walker et al., 2019). The 515F (Parada et al., 2016) and 806R (Apprill et al., 2015) primers were chosen to amplify bacteria and ITS1FI2 (Schmidt et al., 2013) and ITS2 for fungi, as these are the same primers used during amplicon library preparation and sequencing. Cycling conditions on an Agilent AriaMX consisted of 95 C for 3 mins followed by 40 cycles of 95 C for 15 sec, 60 C for 30 sec and 72 C for 30 sec. Following amplification, a melt curve was generated under the following conditions including 95 C for 30 sec, and a melt from 60 C to 90 C increasing in resolution of 0.5 C in increments of a 5 sec soak time. To validate the spike in protocol we selected two sets of fecal samples including 360 samples from a diverse species pool of frogs, lizards, salamanders and snakes and a more targeted approach of 122 fecal samples from three genera of salamanders from the Plethodontidae. (Supplemental Table #). FFecal samples were not weighed in the field, rather, a complete fecal pellet was diluted in an equal volume of sterile water and standardized volume of fecal slurry (250µL) extracted for independent samples.. A volume of 1ul T. halophilus (1874 copies) and 1ul D. bruxellensis (733 copies) were spiked into each fecal sample then DNA was extracted as above, libraries constructed, and amplicon sequenced on an Illumina MiSeq as in (Vargas-Gastélum et al., 2024) .

Summary of the DspikeIn core features:

1.Validation of spiked species: DspikeIn facilitates the verification of OTUs/ASVs originating from spiked species through phylogenetic tree construction (e.g., Bootstrap_phy_tree_with_cophenetic), ensuring accuracy in downstream analyses. 2. Data preprocessing: The package prepares data for analysis by merging multiple OTUs/ASVs associated with spiked species (e.g., Pre_processing_species_list). 3. Scaling factor extraction: It extracts scaling factors from preprocessed data (e.g., calculate_list_average_scaling_factors), providing the foundation for accurate microbial quantification. 4. Conversion to absolute abundance: DspikeIn transforms RA data into absolute counts using established scaling factors (e.g., convert_to_absolute_counts). 5. Bias correction and normalization: The package offers systemic bias correction and dataset normalization through various methods (e.g., normalization_set(ps, method = "tss")). 6. System-specific spiked species retrieval: DspikeIn enables detection of optimal system-specific spiked species retrieval through visualization (regression_plot). 7. Performance assessment: It evaluates the success or failure rates of spiked data retrieval (e.g., calculate_spike_percentage_list and conclusion), helping to guide decisions for current and future sample spiking strategies. 8. Exploration and filtering of taxa: Beyond absolute abundance calculations, DspikeIn offers pathways to explore and filter core and dominant taxa. Features include core microbiome analysis, Random Forest-based selection, and prevalence assessments, all paired with intuitive visualization options to aid interpretation.

Bias correction and normalization methods:

Normalization methods validated in the literature (Bolstad et al., 2003; Robinson and Oshlack, 2010; Abbas-Aghababazadeh et al., 2018) have been selected and modified from several references and packages, including EdgeR, DESeq2, and others. These bias correction methods include:

a) Trimmed Mean of M-values (TMM) Normalization: A scaling normalization method used for RNA-seq count data to account for compositional differences between libraries (Robinson and Oshlack, 2010). b) Cumulative Sum Scaling (CSS): This method assumes equivalent count distributions for low-abundance genes across samples up to a certain threshold. It scales the invariant segment of each sample's count distribution and normalizes counts based on invariant counts across samples. c) Total Sum Scaling (TSS): Converts the feature table into relative abundance by dividing the total reads of each sample. TSS is commonly used for generating relative abundance data. Rarefying: Randomly removes reads from different samples until they all have the same predefined number of reads, ensuring equal library size. This method is considered a standard approach in microbial ecology. d) Centered Log-Ratio Transformation (CLR): Computes log-ratios relative to the geometric mean of all features, used to handle compositional data. e) Counts per Million (CPM): Derived from the LEfSe algorithm, this method is recommended for datasets with very low counts and is often used in differential abundance analysis. f) Relative Log Expression (RLE): Assumes that most features are not differentially expressed and uses relative abundances to calculate the normalization factor. g) RLE is commonly used in RNA-seq normalization. DESeq Normalization: Normalizes counts based on the assumption that most genes are not differentially expressed. DESeq is widely used in RNA-seq data analysis. h) PoissonSeq: A normalization method specifically designed for RNA-seq data, available in the PoissonSeq package. i) Quantile Normalization (QN): Adjusts the distribution of counts so that different samples have the same distribution, used in high-throughput data analysis.

Phylogenetic structure and differential abundance

Microbial phylogenetic diversity was analyzed and compared between D. imitator and D. monticola. Analysis of phylogenetic structure revealed diverse fungal and bacterial communities for both species, reflecting stable microbial associations (Kajihara and Hynson, 2024) (Figures S6a-d, S7a-d). Positive SES values indicate phylogenetic overdispersion driven by competition, while negative values reflect clustering due to environmental filtering favoring related species in similar habitats (Aguirre De Cárcer, 2019; O'Dwyer et al., 2012). Fungi in samples UHM1405.2085_S173 ($p = 0.04$), UHM1419.20860_S114 ($p = 0.04$), and UHM1425.20863_S150 ($p = 0.01$) exhibited environmental filtering. Bacteria, with the exception of UHM437.20864_S162 and UHM1405.20857_S173 samples, exhibited phylogenetic overdispersion. Mean Pairwise Distance (MPD) analysis revealed high similarity between fungal ($p = 0.30$) and bacterial ($p = 0.16$) communities in Desmognathus monticola and Desmognathus imitator, indicating that these species exhibit comparable levels of phylogenetic diversity.

Network topology and stability test

To investigate the ecological role of Basidiobolus in the herptile gut microbiome, we analyzed three absolute abundance-based networks for D. monticola and D. imitator: a complete network, one with network and module hubs removed, and one excluding the Basidiobolus subnetwork. The removal of Basidiobolus had a greater impact on the shape and diameter of network compared to the removal of network and module hubs (Fig. 5d-f, S10a-d, and S11a-d). Metrics such as local efficiency, harmonic centrality, and closeness were similarly affected by the removal of either the Basidiobolus subnetwork or network and module hubs (Fig. 6a-f). Interestingly, the complete network displayed exclusively positive interactions; however, the removal of hubs led to an 11% increase in negative interactions, while removing the Basidiobolus subnetwork resulted in a 14.4% increase. Connectivity and modularity networks emphasized the role of individual taxa in maintaining network integrity, with Basidiobolus specifically identified as a connector taxon facilitating inter-module interactions (Fig. 4a, 5d-f, and S10a-c).

Key metrics of network robustness, including the largest connected component (LCC) and modularity, showed minimal changes after removing the Basidiobolus subnetwork, with a moderate shift in average path length (Fig. 6a-i). The fraction size of the LCC versus the fraction of excluded nodes exhibited a similar smooth decreasing trend in both scenarios: removal of network and module hubs and removal of the Basidiobolus subnetwork (Fig. 6d). High redundancy indicates a robust network, as overlapping connections, providing functional buffering. Removing low-redundancy species from the network caused a ~38% reduction in LCC size (from 308 to 191), highlighting their crucial role in maintaining network connectivity, a key factor in overall network stability. In the complete network, species displayed varying redundancy levels, with a mean of 1.2 and a median of 1.0, reflecting the network's robustness. Nodes such as OTU256:Alistipes inops and OTU268 occupied key positions in the top-right quadrant, signifying their roles as both highly connected and locally stable hubs. The removal of network and module hubs reduced connectivity, redistributing nodes to lower degree and redundancy regions. Plotting nodes based on connectivity metrics, such as betweenness and degree, revealed that Basidiobolus (OTU69) exhibited both high betweenness and degree, underscoring its role as a central hub and bridge within the network (Fig. 7a-c). Basidiobolus (OTU69) was positioned in the low redundancy, high degree, and efficiency region, indicating it may play a less critical role in stabilizing the network but serves as a key bridge among communities. In the complete network, the spatial position of Basidiobolus (OTU69) on the betweenness versus degree plot highlighted its structural importance as a bridge or broker. Its high effective size underscores its unique and essential role within the network.

Core microbiome dynamics

Rikenella was unique to the Anuran clade in the Mississippi Alluvial Plain, correlating with carnivorous hosts with metamorphic physiology. Lactococcus was widespread across diverse hosts and habitats, while Erysipelatoclostridium occurred in oviparous and insectivorous hosts.

Eurycea exhibited a more balanced fungal community structure, with notable contributions from Mortierella and Backusella based on AA. Host- and region-specific patterns in the abundance of Basidiobolus and Mortierella were evident in AA but obscured in RA analyses.

OUT vs ASV and GCN correction debate

The debate over OTUs versus ASVs [78, 79] parallels discussions on gene copy number (GCN) correction for the 16S rRNA marker. While proponents argue that GCN correction enhances accuracy [75, 118], critics highlight its limitations [55, 119]. Currently, GCN correction is not integrated into the DspikeIn package, it can be applied to RA data prior to conversion to absolute counts using tools such as the q2-gcn-norm plugin in Qiime2 (rrnDB v5.7) [23] or methods outlined by Louca et al. [55] including PICRUSt, CopyRighter, and PAPRICA. However, due to the inherent variability in rDNA gene copy numbers [56], GCN correction for ITS data is generally not recommended, except when applied to specific taxa with well-characterized loci to enable accurate normalization of abundance.

Spike-in volume Protocol,

The species Tetragenococcus halophilus (bacterial spike; ATCC33315) and Dekkera bruxellensis (fungal spike; WLP4642-White Labs) were selected as taxa to spike into gut microbiome samples as they were not found in an extensive collection of wildlife skin (GenBank BioProjects: PRJNA1114724, PRJNA 1114659) or gut microbiome samples (PRJNA##this study##). Stock cell suspensions of both microbes were grown in either static tryptic soy broth (T. halophilus) or potato dextrose broth (D. bruxellensis) for 72 hours then serially diluted and optical density (OD600) determined on a ClarioStar plate reader. Cell suspensions with an optical density of 1.0, 0.1, 0.01, 0.001 were DNA extracted using the Qiagen DNeasy Powersoil Pro Kit. These DNA isolations were used as standards to determine the proper spike in volume of cells to represent 0.1-10% of a sample (Rao et al., 2021b) Fecal pellets (3.1 ± 1.6 mg; range = 1 – 5.1 mg) from an ongoing live animal study using wood frogs (Lithobates sylvaticus) were used to standardize the input material for the development of this protocol. A total of (n=9) samples were used to validate the spike in protocol. Each fecal sample was homogenized in 1mL of sterile molecular grade water then 250uL of fecal slurry was DNA extracted as above with and without spiked cells.

Two approaches were used to evaluate the target spike-in of 0.1-10%, the range of effective spike-in percentage described in (Rao et al., 2021b), including 1) an expected increase of qPCR cycle threshold (Ct) value that is proportional to the amount of spiked cells and 2) the expected increase in copy number of T. halophilus and D. bruxellensis in spiked vs. unspiked samples. A standard curve was generated using a synthetic fragment of DNA for the 16S-V4 rRNA and ITS1 rDNA regions of T. halophilus and D. bruxellensis, respectively. The standard curve was used to convert

Ct values into log copy number for statistical analyses (detailed approach in[2, 3]) using the formula y = -0.2426x + 10.584 for T. halophilus and y = -0.3071x + 10.349 for D. bruxellensis, where x is the average Ct for each unknown sample.

Quantitative PCR (qPCR) was used to compare known copy numbers from synthetic DNA sequences of T. halophilus and D. bruxellensis to DNA extractions of T. halophilus and D. bruxellensis independently, and wood frog fecal samples with and without spiked cells. SYBR qPCR assays were run at 20ul total volume including 10ul 2X Quantabio PerfeCTa SYBR Green Fastmix, 1ul of 10uM forward and reverse primers, 1ul of ArcticEnzymes dsDNAse master mix clean up kit, and either 1ul of DNA for D. bruxellensis or 3ul for T. halophilus. Different volumes of DNA were chosen for amplification of bacteria and fungi due to previous optimization of library preparation and sequencing steps [3]. The 515F [4] and 806R [5] primers were chosen to amplify bacteria and ITS1FI2 [6] and ITS2 for fungi, as these are the same primers used during amplicon library preparation and sequencing. Cycling conditions on an Agilent AriaMX consisted of 95 C for 3 mins followed by 40 cycles of 95 C for 15 sec, 60 C for 30 sec and 72 C for 30 sec. Following amplification, a melt curve was generated under the following conditions including 95 C for 30 sec, and a melt from 60 C to 90 C increasing in resolution of 0.5 C in increments of a 5 sec soak time. To validate the spike in protocol we selected two sets of fecal samples including 360 samples from a diverse species pool of frogs, lizards, salamanders and snakes and a more targeted approach of 122 fecal samples from three genera of salamanders from the Plethodontidae. (Supplemental Table #). FFecal samples were not weighed in the field, rather, a complete fecal pellet was diluted in an equal volume of sterile water and standardized volume of fecal slurry (250µL) extracted for independent samples.. A volume of 1ul T. halophilus (1874 copies) and 1ul D. bruxellensis (733 copies) were spiked into each fecal sample then DNA was extracted as above, libraries constructed, and amplicon sequenced on an Illumina MiSeq as in [7] .

Summary of the DspikeIn core features:

1.Validation of spiked species: DspikeIn facilitates the verification of OTUs/ASVs originating from spiked species through phylogenetic tree construction (e.g., Bootstrap_phy_tree_with_cophenetic), ensuring accuracy in downstream analyses. 2. Data preprocessing: The package prepares data for analysis by merging multiple OTUs/ASVs associated with spiked species (e.g., Pre_processing_species_list). 3. Scaling factor extraction: It extracts scaling factors from preprocessed data (e.g., calculate_list_average_scaling_factors), providing the foundation for accurate microbial quantification. 4. Conversion to absolute abundance: DspikeIn transforms RA data into absolute counts using established scaling factors (e.g., convert_to_absolute_counts). 5. Bias correction and normalization: The package offers systemic bias correction and dataset normalization through various methods (e.g., normalization_set(ps, method = "tss")). 6. System-specific spiked species retrieval: DspikeIn enables detection of optimal system-specific spiked species retrieval through visualization (regression_plot). 7. Performance assessment: It evaluates the success or failure rates of spiked data retrieval (e.g., calculate_spike_percentage_list and conclusion), helping to guide decisions for current and future sample spiking strategies. 8. Exploration and filtering of taxa: Beyond absolute abundance calculations, DspikeIn offers pathways to explore and filter core and dominant taxa. Features include core microbiome analysis, Random Forest-based selection, and prevalence assessments, all paired with intuitive visualization options to aid interpretation.

Bias correction and normalization methods:

Normalization methods validated in the literature (Bolstad et al., 2003; Robinson and Oshlack, 2010; Abbas-Aghababazadeh et al., 2018) have been selected and modified from several references and packages, including EdgeR, DESeq2, and others. These bias correction methods include:

a) Trimmed Mean of M-values (TMM) Normalization: A scaling normalization method used for RNA-seq count data to account for compositional differences between libraries (Robinson and Oshlack, 2010). b) Cumulative Sum Scaling (CSS): This method assumes equivalent count distributions for low-abundance genes across samples up to a certain threshold. It scales the invariant segment of each sample's count distribution and normalizes counts based on invariant counts across samples. c) Total Sum Scaling (TSS): Converts the feature table into relative abundance by dividing the total reads of each sample. TSS is commonly used for generating relative abundance data. Rarefying: Randomly removes reads from different samples until they all have the same predefined number of reads, ensuring equal library size. This method is considered a standard approach in microbial ecology. d) Centered Log-Ratio Transformation (CLR): Computes log-ratios relative to the geometric mean of all features, used to handle compositional data. e) Counts per Million (CPM): Derived from the LEfSe algorithm, this method is recommended for datasets with very low counts and is often used in differential abundance analysis. f) Relative Log Expression (RLE): Assumes that most features are not differentially expressed and uses relative abundances to calculate the normalization factor. g) RLE is commonly used in RNA-seq normalization. DESeq Normalization: Normalizes counts based on the assumption that most genes are not differentially expressed. DESeq is widely used in RNA-seq data analysis. h) PoissonSeq: A normalization method specifically designed for RNA-seq data, available in the PoissonSeq package. i) Quantile Normalization (QN): Adjusts the distribution of counts so that different samples have the same distribution, used in high-throughput data analysis.

#### Phylogenetic structure and differential abundance

Microbial phylogenetic diversity was analyzed and compared between D. imitator and D. monticola. Analysis of phylogenetic structure revealed diverse fungal and bacterial communities for both species, reflecting stable microbial associations (Kajihara and Hynson, 2024) (Figures S6a-d, S7a-d). Positive SES values indicate phylogenetic overdispersion driven by competition, while negative values reflect clustering due to environmental filtering favoring related species in similar habitats [9, 10]. Fungi in samples UHM1405.2085_S173 ($p = 0.04$), UHM1419.20860_S114 ($p = 0.04$), and UHM1425.20863_S150 ($p = 0.01$) exhibited environmental filtering. Bacteria, with the exception of UHM437.20864_S162 and UHM1405.20857_S173 samples, exhibited phylogenetic overdispersion. Mean Pairwise Distance (MPD) analysis revealed high similarity between fungal ($p = 0.30$) and bacterial ($p = 0.16$) communities in Desmognathus monticola and Desmognathus imitator, indicating that these species exhibit comparable levels of phylogenetic diversity.

#### Network topology and stability test

To investigate the ecological role of Basidiobolus in the herptile gut microbiome, we analyzed three absolute abundance-based networks for D. monticola and D. imitator: a complete network, one with network and module hubs removed, and one excluding the Basidiobolus subnetwork. The removal of Basidiobolus had a greater impact on the shape and diameter of network compared to the removal of network and module hubs (Fig. 5d-f, S10a-d, and S11a-d). Metrics such as local efficiency, harmonic centrality, and closeness were similarly affected by the removal of either the Basidiobolus subnetwork or network and module hubs (Fig. 6a-f). Interestingly, the complete network displayed exclusively positive interactions; however, the removal of hubs led to an 11% increase in negative interactions, while removing the Basidiobolus subnetwork resulted in a 14.4% increase. Connectivity and modularity networks emphasized the role of individual taxa in maintaining network integrity, with Basidiobolus specifically identified as a connector taxon facilitating inter-module interactions (Fig. 4a, 5d-f, and S10a-c).

Key metrics of network robustness, including the largest connected component (LCC) and modularity, showed minimal changes after removing the Basidiobolus subnetwork, with a moderate shift in average path length  (Fig. 6a-i). The fraction size of the LCC versus the fraction of excluded nodes exhibited a similar smooth decreasing trend in both scenarios: removal of network and module hubs and removal of the Basidiobolus subnetwork (Fig. 6d). High redundancy indicates a robust network, as overlapping connections, providing functional buffering. Removing low-redundancy species from the network caused a ~38% reduction in LCC size (from 308 to 191), highlighting their crucial role in maintaining network connectivity, a key factor in overall network stability. In the complete network, species displayed varying redundancy levels, with a mean of 1.2 and a median of 1.0, reflecting the network's robustness. Nodes such as OTU256:Alistipes inops and OTU268 occupied key positions in the top-right quadrant, signifying their roles as both highly connected and locally stable hubs. The removal of network and module hubs reduced connectivity, redistributing nodes to lower degree and redundancy regions. Plotting nodes based on connectivity metrics, such as betweenness and degree, revealed that Basidiobolus (OTU69) exhibited both high betweenness and degree, underscoring its role as a central hub and bridge within the network (Fig. 7a-c). Basidiobolus (OTU69) was positioned in the low redundancy, high degree, and efficiency region, indicating it may play a less critical role in stabilizing the network but serves as a key bridge among communities. In the complete network, the spatial position of Basidiobolus (OTU69) on the betweenness versus degree plot highlighted its structural importance as a bridge or broker. Its high effective size underscores its unique and essential role within the network.

Core microbiome dynamics

Rikenella was unique to the Anuran clade in the Mississippi Alluvial Plain, correlating with carnivorous hosts with metamorphic physiology. Lactococcus was widespread across diverse hosts and habitats, while Erysipelatoclostridium occurred in oviparous and insectivorous hosts.

Eurycea exhibited a more balanced fungal community structure, with notable contributions from Mortierella and Backusella based on AA. Host- and region-specific patterns in the abundance of Basidiobolus and Mortierella were evident in AA but obscured in RA analyses.

OUT vs ASV and GCN correction debate

The debate over OTUs versus ASVs [11, 12] parallels discussions on gene copy number (GCN) correction for the 16S rRNA marker. While proponents argue that GCN correction enhances accuracy [13, 14], critics highlight its limitations [15, 16]. Currently, GCN correction is not integrated into the DspikeIn package, it can be applied to RA data prior to conversion to absolute counts using tools such as the q2-gcn-norm plugin in Qiime2 (rrnDB v5.7) [17] or methods outlined by Louca et al. [15] including PICRUSt, CopyRighter, and PAPRICA. However, due to the inherent variability in rDNA gene copy numbers [18], GCN correction for ITS data is generally not recommended, except when applied to specific taxa with well-characterized loci to enable accurate normalization of abundance.


1. Rao C, Coyte KZ, Bainter W, Geha RS, Martin CR, Rakoff-Nahoum S. Multi-kingdom ecological drivers of microbiota assembly in preterm infants. Nature 2021; 591: 633–638.

2. Romer AS, Grinath JB, Moe KC, Walker DM. Host microbiome responses to the Snake Fungal Disease pathogen (Ophidiomyces ophidiicola) are driven by changes in microbial richness. Sci Rep 2022; 12.

9. Aguirre De Cárcer D. A conceptual framework for the phylogenetically constrained assembly of microbial communities. Microbiome 2019; 7.

10. O'Dwyer JP, Kembel SW, Green JL. Phylogenetic Diversity Theory Sheds Light on the Structure of Microbial Communities. PLoS Comput Biol 2012; 8.

11. Callahan BJ, McMurdie PJ, Holmes SP. Exact sequence variants should replace operational taxonomic units in marker-gene data analysis. ISME Journal 2017; 11: 2639–2643.

12. Chiarello M, McCauley M, Villéger S, Jackson CR. Ranking the biases: The choice of OTUs vs. ASVs in 16S rRNA amplicon data analysis has stronger effects on diversity measures than rarefaction and OTU identity threshold. PLoS One 2022; 17.

13. Pollock J, Glendinning L, Wisedchanwet T, Watson M. The madness of microbiome: Attempting to find consensus 'best practice' for 16S microbiome studies. Appl Environ Microbiol . 2018. American Society for Microbiology. , 84

14. Starke R, Pylro VS, Morais DK. 16S rRNA Gene Copy Number Normalization Does Not Provide More Reliable Conclusions in Metataxonomic Surveys. Microb Ecol 2021; 81: 535–539.

15. Louca S, Doebeli M, Parfrey LW. Correcting for 16S rRNA gene copy numbers in microbiome surveys remains an unsolved problem. Microbiome 2018; 6.

16. Gao Y, Wu M. Accounting for 16S rRNA copy number prediction uncertainty and its implications in bacterial diversity analyses. ISME Communications 2023; 3.

17. Bolyen E, RJR, DMR, BNA, ACC, A-GGA, AH, AEJ, AM, AF and BY. Reproducible, interactive, scalable and extensible microbiome data science using QIIME 2. Nat Biotechnol . 2019. Nature Publishing Group. , 37: 850–852

18. Lavrinienko A, Jernfors T, Koskimäki JJ, Pirttilä AM, Watts PC. Does Intraspecific Variation in rDNA Copy Number Affect Analysis of Microbial Communities? Trends Microbiol . 2021. Elsevier Ltd. , 29: 19–27

Supplemental Figures and Tables

Table S1 Terminology of metrics used to explain gut microbial associations (Coyte et al., n.d.; Deng et al., 2012; Ghotbi et al., 2025; Kajihara and Hynson, 2024; Van Der Heijden and Hartmann, 2016)

| **Term** | **Definition** |
|---|---|
| AmongModuleConnectivity | The degree to which nodes connect across different groups or modules within the network. |
| WithinModuleConnectivity | The extent to which nodes connect within the same group or module. |
| Average path length | The average path length is the mean of the shortest path lengths between all pairs of nodes in a network. |
| Betweenness Centrality | A measure of how often a node lies on the shortest path between other nodes. |
| Closeness Centrality | Describes how quickly a node can interact with all other nodes. |
| Degree | The number of direct connections (edges) a node has to other nodes. |
| Edge Density | The proportion of possible edges that exist in the network |
| Global Efficiency | The average inverse shortest path length across all node pairs in the network. |
| Local Efficiency | Efficiency within the neighborhood of a node, reflecting the fault tolerance of the network at a local level. |
| Module Hubs | A group of nodes that are more interconnected with each other than with nodes outside the group. |
| Network Hubs | Key nodes that have a disproportionately high number of connections to other nodes. |
| Subnetwork | A smaller network that is part of a larger one, focusing on a specific subset of nodes and their connections. |
| Eigenvector Centrality | Measures the influence of a node based not just on its direct connections but also on the importance of the nodes it connects to. |
| Harmonic Centrality | A variation of closeness centrality that accounts for the distance of each node from others, favoring nodes that are closer to more nodes. |
| Coreness | Indicates how central a node is in the network |
| Authority Score | A measure of how important a node is within the network, based on the number of connections from important nodes pointing to it. |
| Transitivity | Quantifies how well nodes in a network cluster together. |
| Redundancy Ratio | The proportion of duplicated connections. Can be measured the degree of overlap in connections between nodes |
| Clustering Coefficient | Measures the degree to which nodes in a graph tend to cluster together. |
| Largest Connected Component | The size of the largest connected subgraph. |
| Constraint | Measures how much a node is constrained by its neighbors, often due to redundancy in connections |

A B

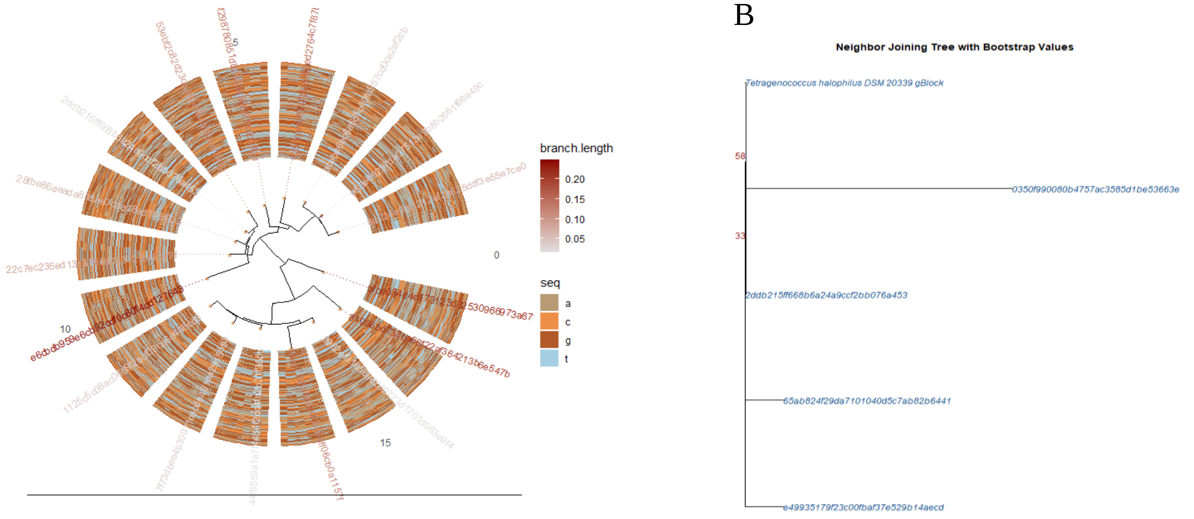


C

| Sample | Total_Reads | Spiked_Total_Reads | Percentage | Result |
|---|---|---|---|---|
| spiked.blank.20433_S84 | 8 | 8 | 100.000000 | failed |
| spiked.blank.20817_S84 | 46,822 | 46,801 | 99.955149 | failed |
| Std2uL.20625_S84 | 62,463 | 62,441 | 99.964779 | failed |
| StdSwab1uL.20624_S72 | 25,008 | 17,809 | 71.213212 | failed |
| UHM369.20773_S31 | 7,498 | 234 | 3.120832 | passed |
| UHM370.20774_S43 | 3,598 | 86 | 2.390217 | passed |
| UHM372.20775_S55 | 253 | 30 | 11.857708 | passed |
| UHM373.20776_S67 | 3,156 | 84 | 2.661597 | passed |
| UHM374.20777_S79 | 4,280 | 70 | 1.635514 | passed |
| UHM375.20778_S91 | 6,872 | 709 | 10.317229 | passed |

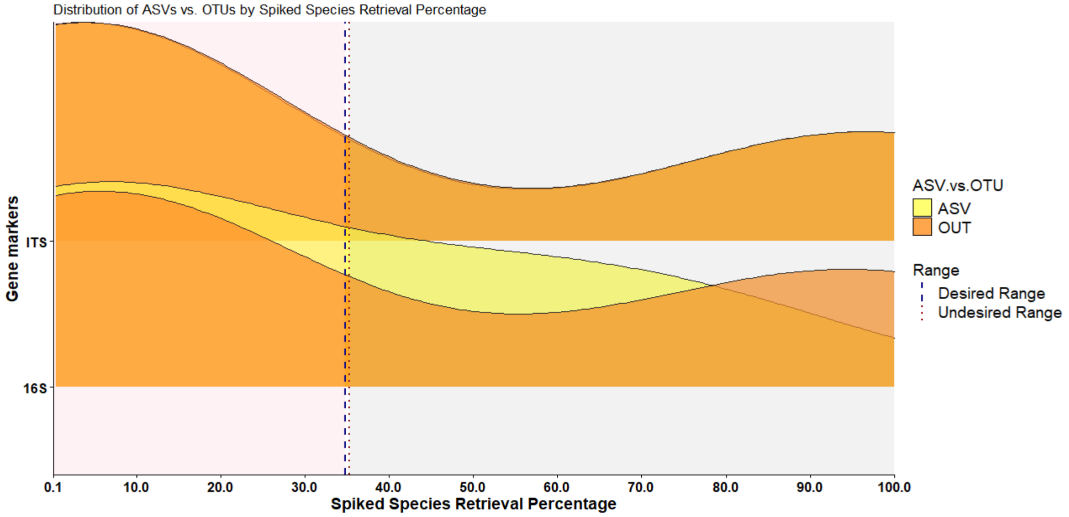


Figure S1 presents a phylogenetic analysis of Amplicon Sequence Variants (ASVs) derived from spiked species. Panel (A) shows the phylogenetic tree illustrating the distances of ASVs from spiked species. Panel (B) compares the phylogenetic distances between positive controls and ASVs rooted in the spiked species present in the samples. Panel (C) evaluates the accuracy of ASV and Operational Taxonomic Unit (OTU) approaches in retrieving spiked species, with retrieval quantified using the calculate_spike_percentage function embedded in DspikeIn.

ITS

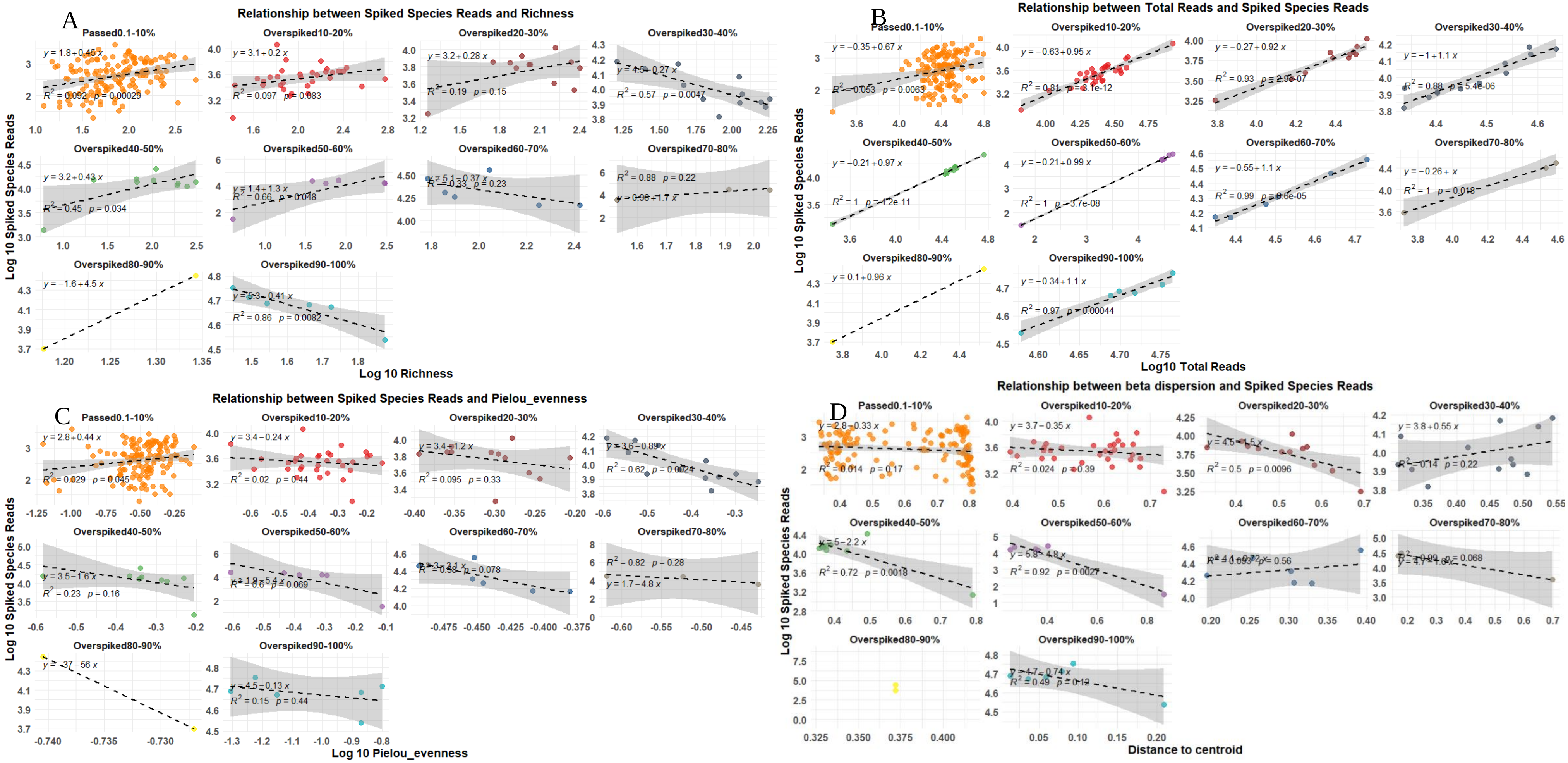

Figure S2 shows the linear relationships between spiked species reads and biological indices in the fungal dataset. Panel (A) presents the relationship with richness, while panel (B) depicts the relationship between the abundance of spiked species and the total abundance. Variations in evenness are shown in panel (C), and panel (D) illustrates the variations in distance to centroid with increasing retrieval ranges of spiked species.

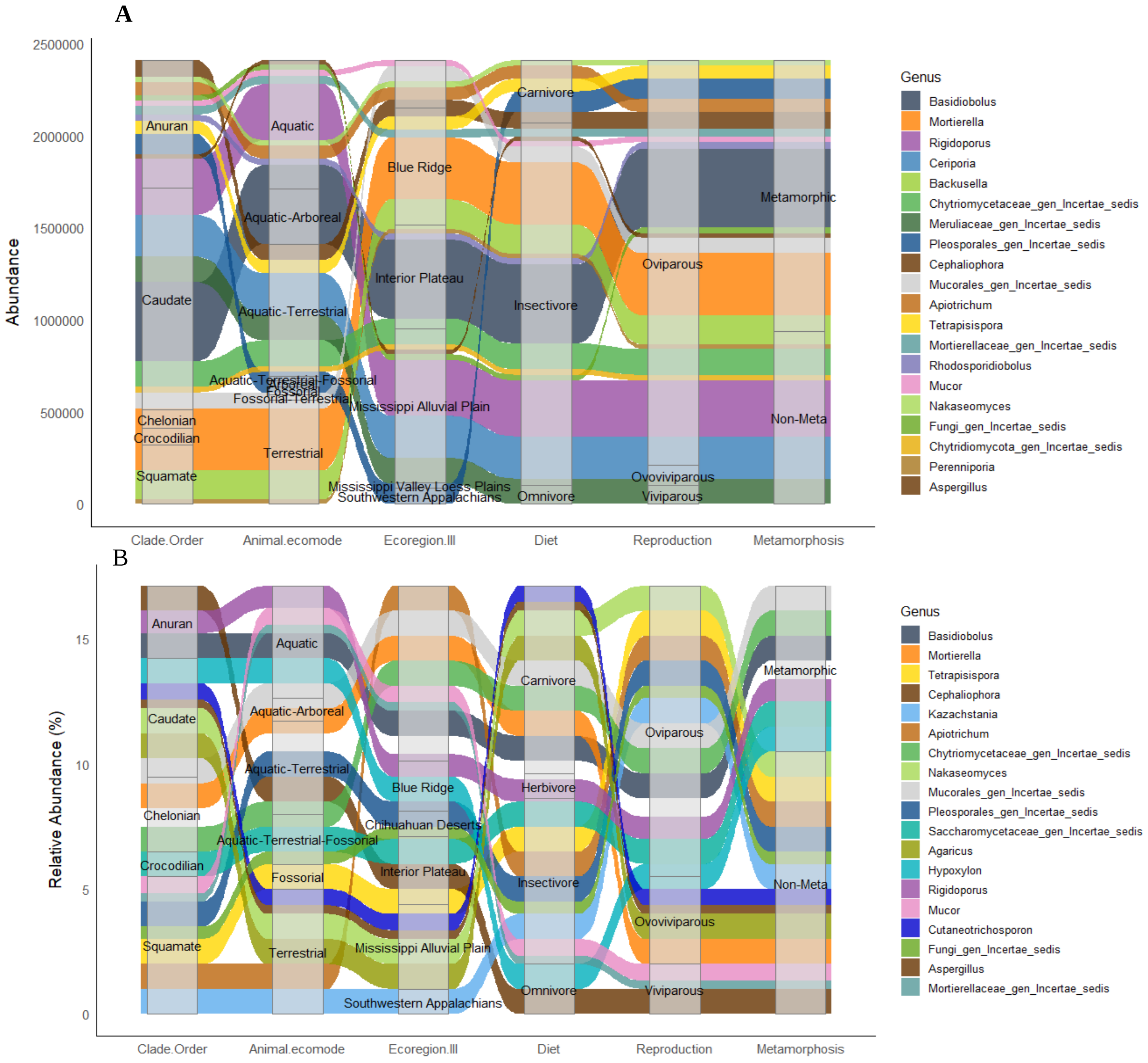


Figure S3 Fungal genus distribution across herptile clade orders, shown by absolute abundance (A) and relative abundance (B) in alluvial plots. Plots illustrate the flow of shared bacteria across ecological or life history, including host clade order, ecomode, ecoregion, diet, reproduction type, and metamorphosis status. Similar color-code was used for genus in both plots for easy traceability.

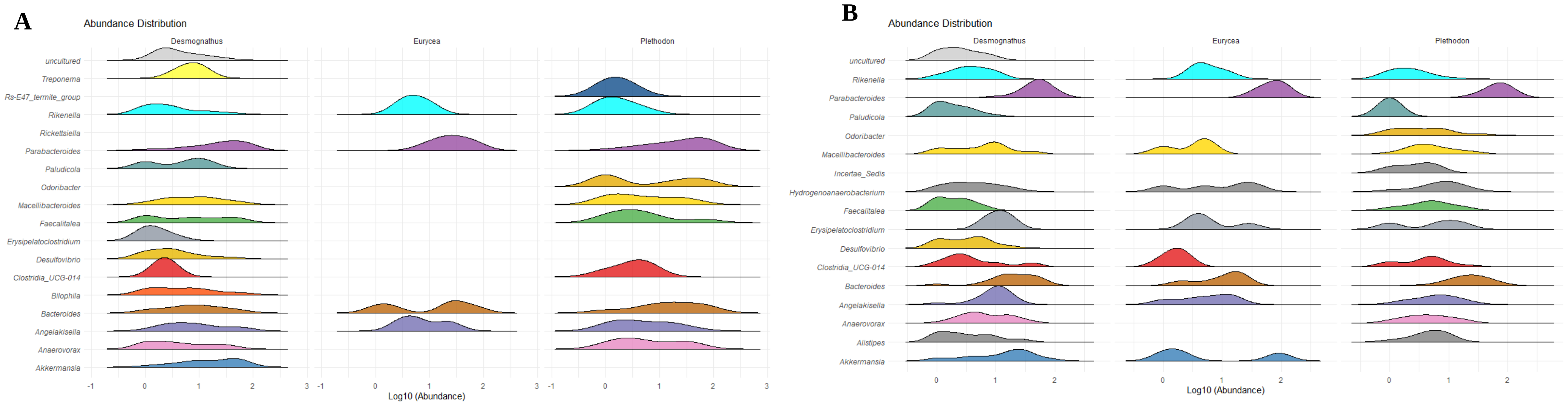


Figure S4 $Log_{10}$ proportion distribution of Random Forest-selected host-associated bacterial OTUs among the *Desmognathus*, *Plethodon*, and *Eurycea* genera, based on absolute (A) versus relative (B) abundance.

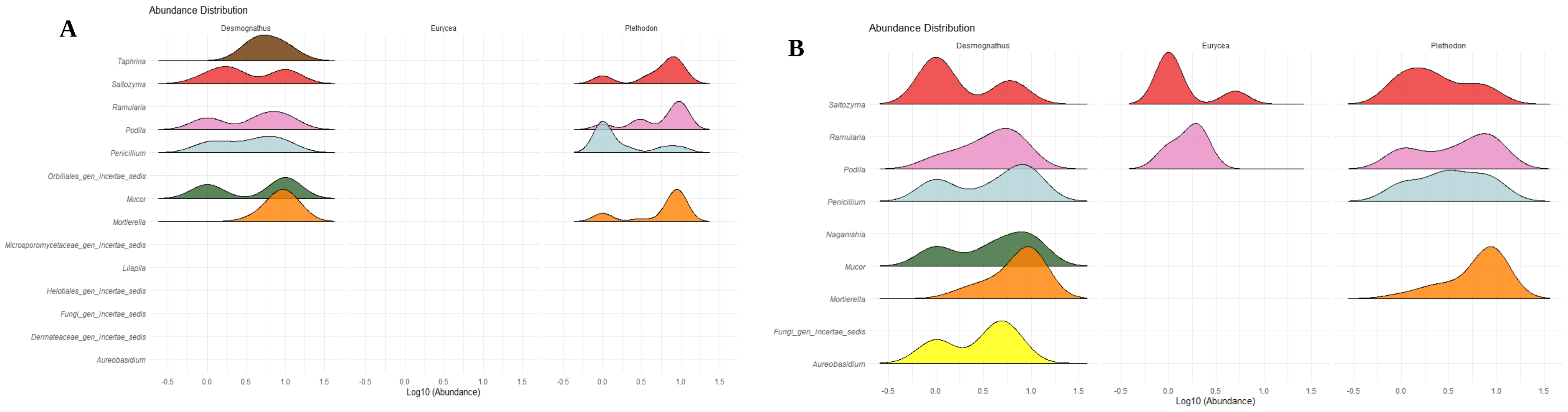


Figure S5 $Log_{10}$ proportion distribution of Random Forest-selected host-associated fungal OTUs among the *Desmognathus*, *Plethodon*, and *Eurycea* genera, based on absolute (A) versus relative (B) abundance.

.

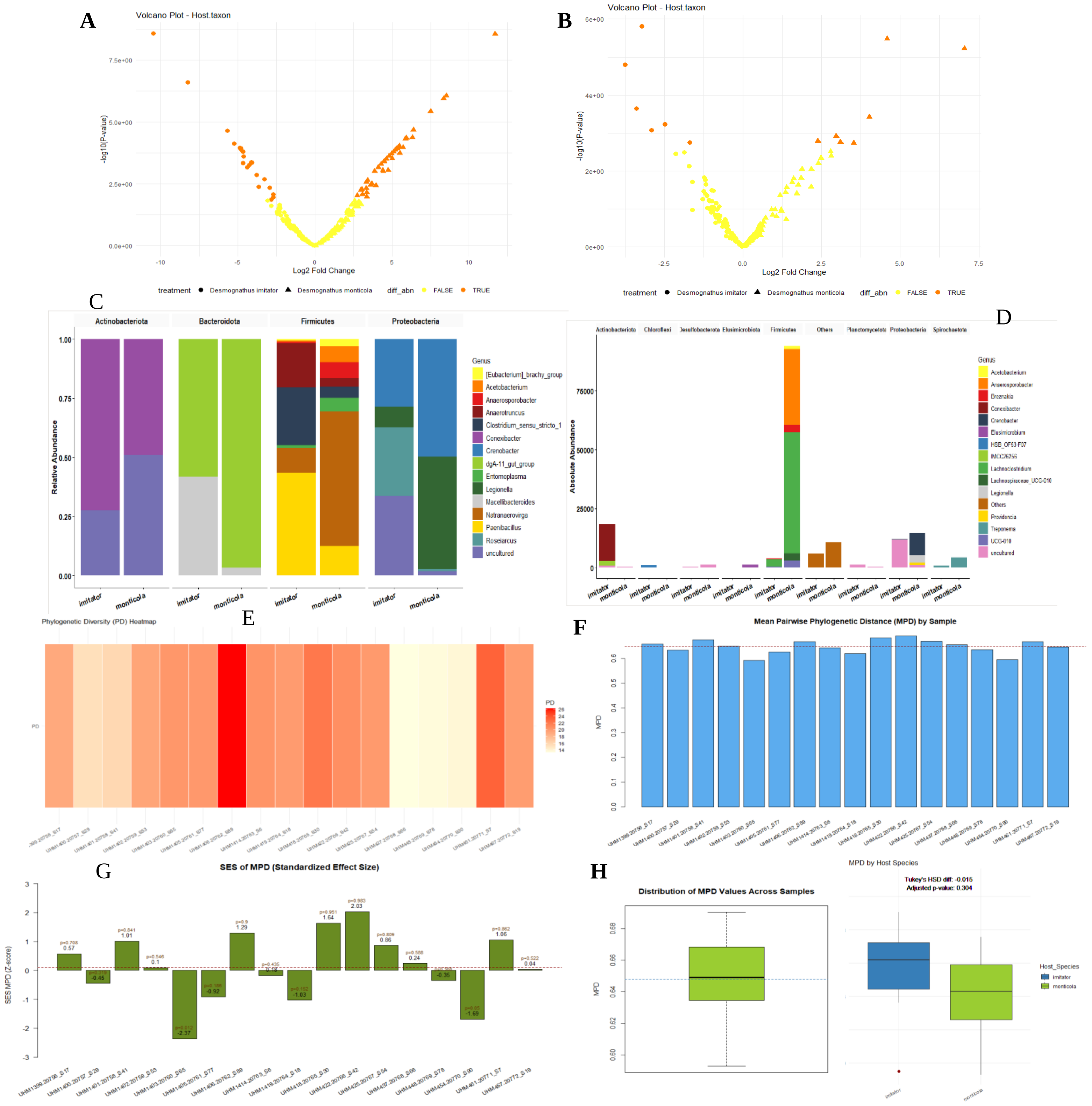


Figure S6 Volcano plots displaying significant OTUs (orange) and nonsignificant OTUs (yellow) for *D. monticola* and *D. imitator*, based on differential abundance analysis using the edgeR method in DspikeIn for absolute (A) and relative counts (B). Counts were normalized with the trimmed mean of M-values (TMM) prior to analysis. The edgeR method (embedded in DspikeIn) uses a generalized linear model (GLM), with statistical significance evaluated via a quasi-likelihood F-test (QLF) to account for biological variability and overdispersion. Significantly enriched taxa are plotted for absolute (C) and relative abundance data (D). Phylogenetic diversity and mean phylogenetic distance are shown (E-H), including comparisons of mean phylogenetic distance between OTUs (E) and within samples (F). Standardized effect size comparisons of phylogenetic distances (G) and mean pairwise phylogenetic distances (H) are plotted for hosts *D. monticola* and *D. imitator*.

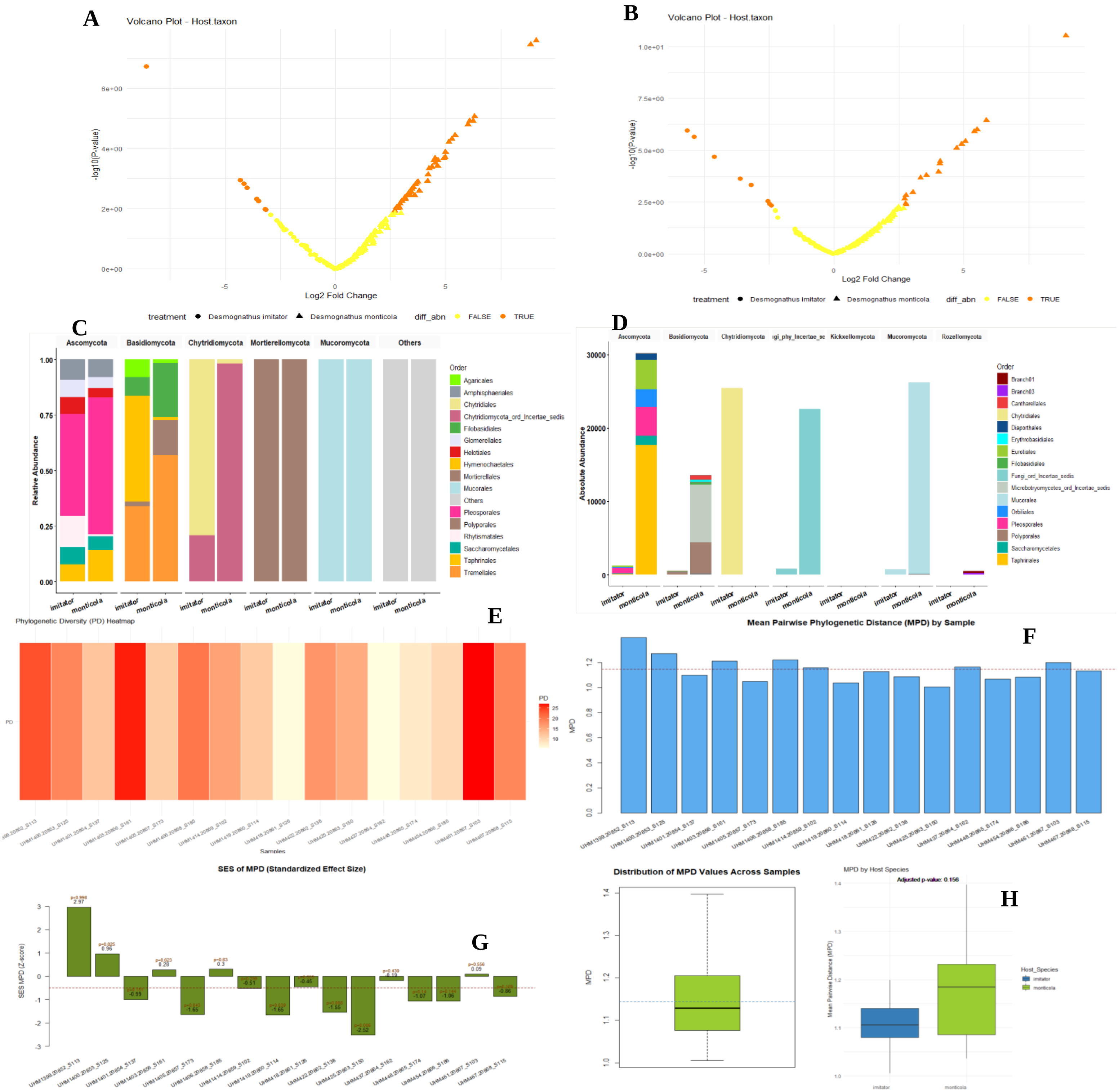

Figure S7 Volcano plots displaying significant OTUs (orange) and nonsignificant OTUs (yellow) for *D. monticola* and *D. imitator*, based on differential abundance analysis using the edgeR method in DspikeIn for absolute (A) and relative counts (B). Counts were normalized with the trimmed mean of M-values (TMM) prior to analysis. The edgeR method (embedded in DspikeIn) uses a generalized linear model (GLM), with statistical significance evaluated via a quasi-likelihood F-test (QLF) to account for biological variability and overdispersion. Significantly enriched taxa are plotted for absolute (C) and relative abundance data (D). Phylogenetic diversity and mean phylogenetic distance are shown (E-H), including comparisons of mean phylogenetic distance between OTUs (E) and within samples (F). Standardized effect size comparisons of phylogenetic distances (G) and mean pairwise phylogenetic distances (H) are plotted for hosts *D. monticola* and *D. imitator*.

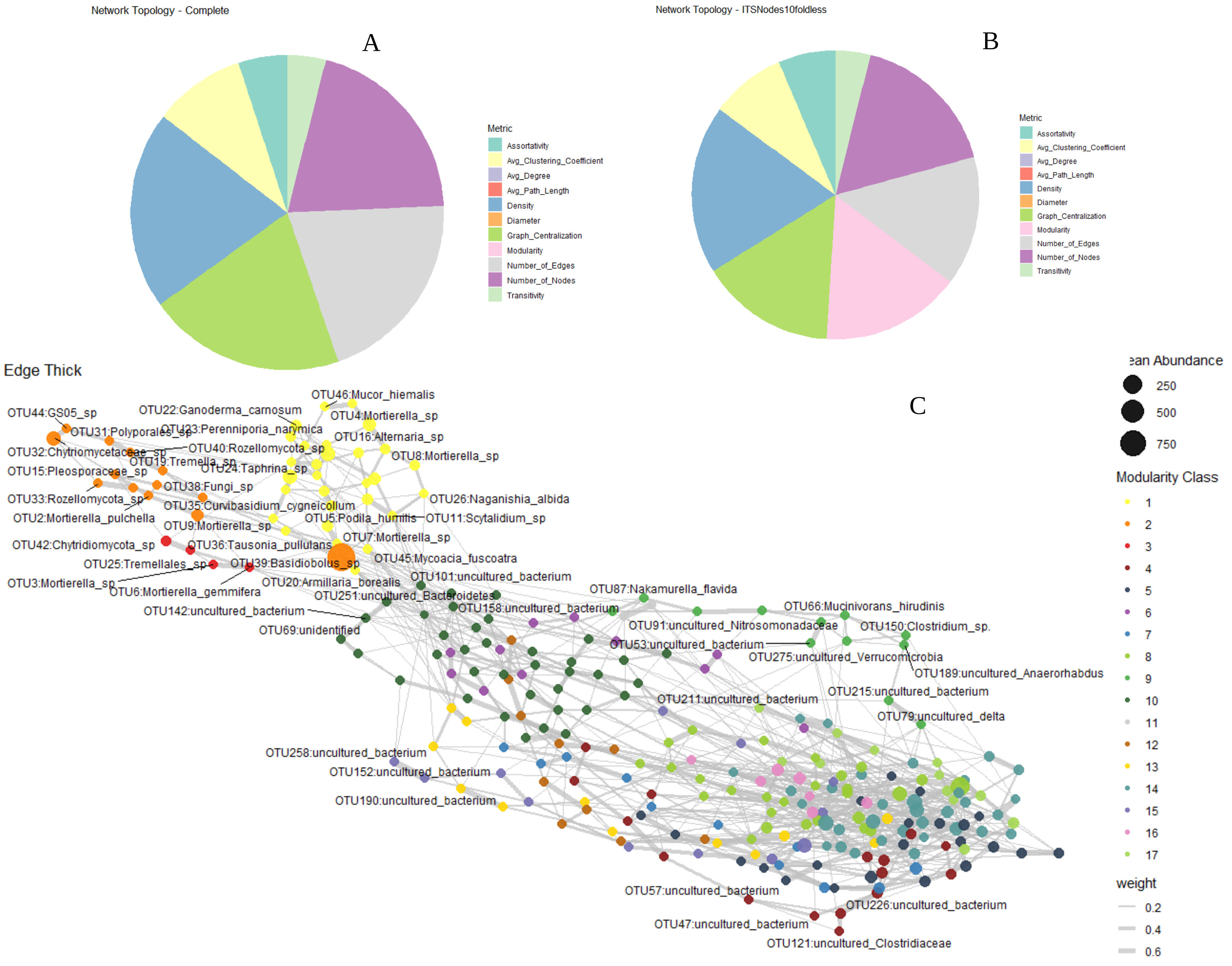


Figure S8 Comparison of network topographic properties across different conditions: (A) complete network, (B) complete network with copy number correction, achieved by dividing all fungal abundances by 10 (C) modularity of cross-domain network associations constructed using absolute counts for the complete network with copy number correction (dividing all fungal abundances by 10) from the gut microbiome of Dendrobates imitator and Dendrobates monticola species. The networks exhibit an organic structure, with edge weight represented by edge thickness, node sizes are representative of mean abundance, and phyla color-coded.

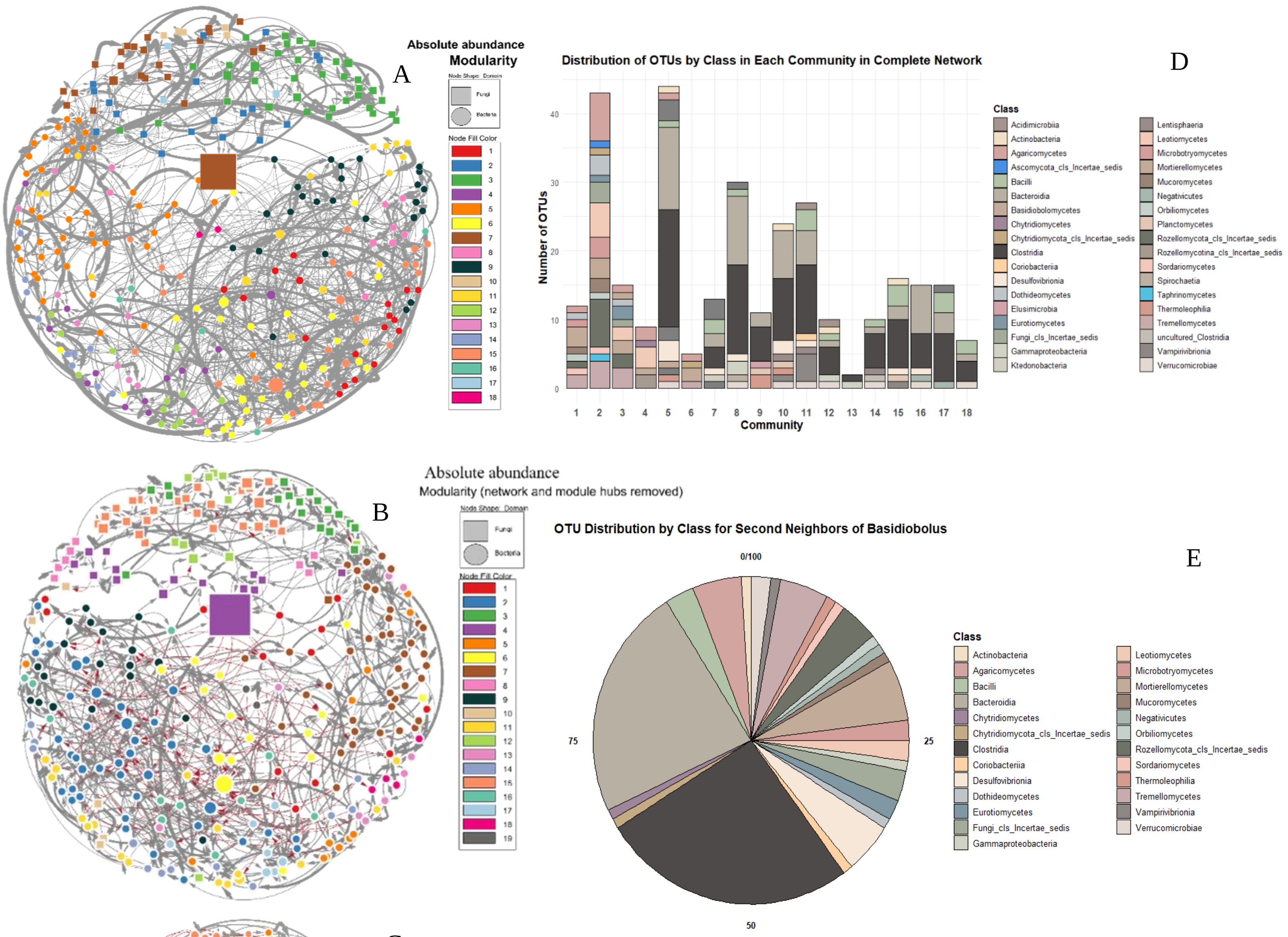


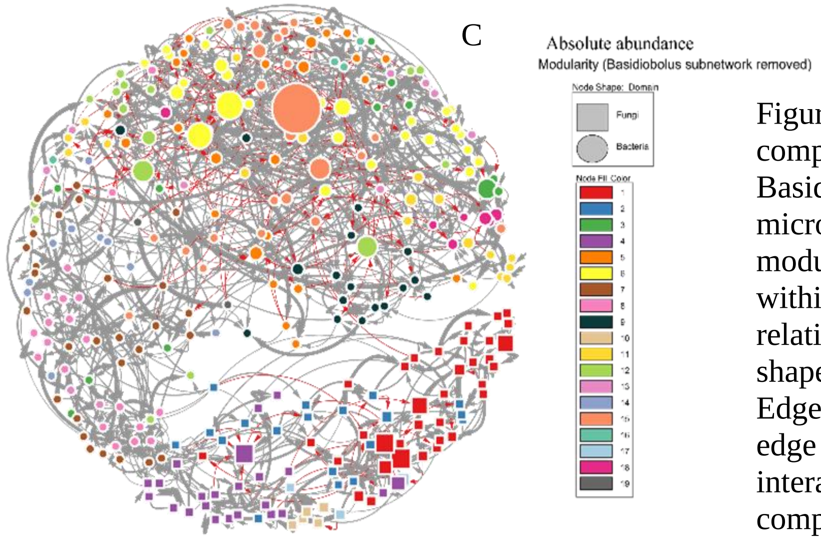


Figure S9 Illustrates the modularity of networks for complete (A), network and module hubs removed (B), and Basidiobolus subnetwork removed (C) from the gut microbiomes of *D. monticola* and *D. imitator*. In these module networks, nodes of the same color represent taxa within the same module. Node size corresponds to the relativized absolute abundance of each taxon, while node shapes distinguish bacteria (circles) from fungi (squares). Edge thickness represents the degree of connections, and edge colors indicate positive (gray) or negative (red) interactions. The module/community members of the complete network are shown in the bar plot (D). Basidiobolus' second neighbors and the number of OTUs for each member are presented in panel (E).v

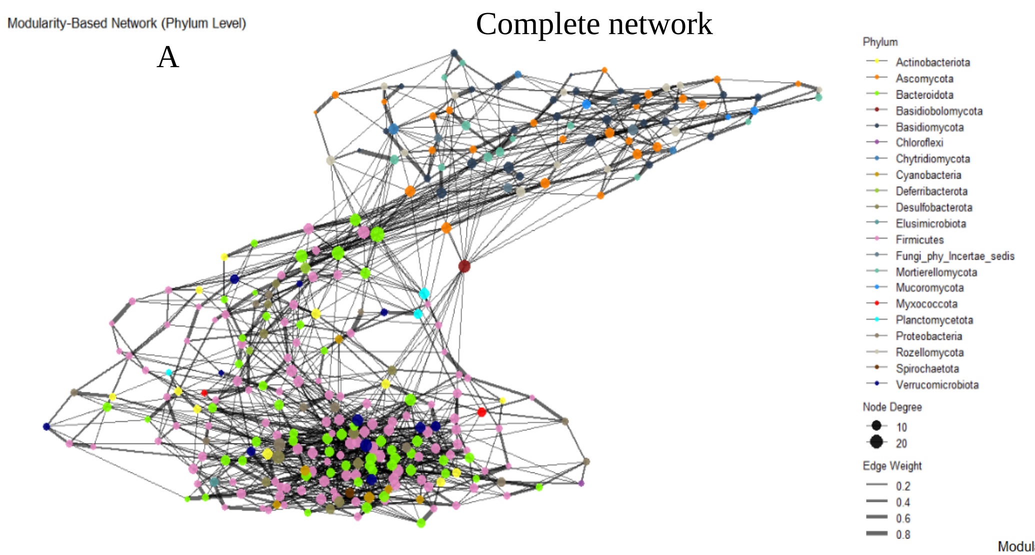


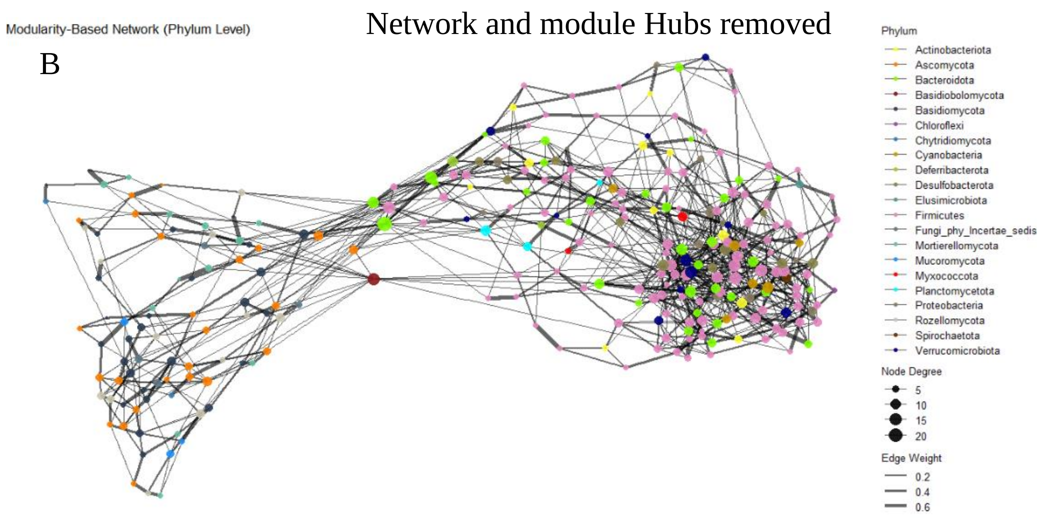


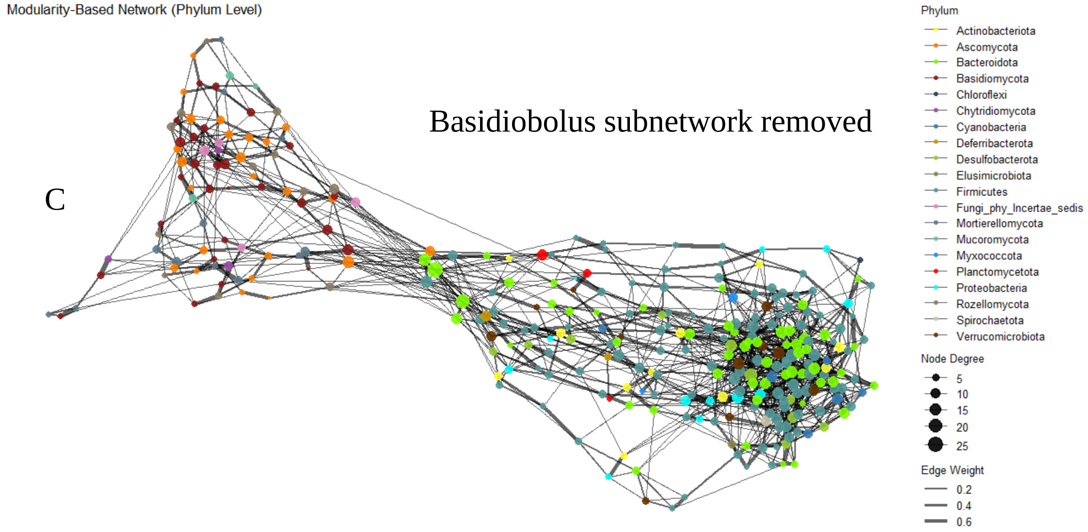


Figure S10 shows the modularity of cross-domain network associations constructed using absolute counts for the complete network (A), the network with module hubs removed (B), and the network with the Basidiobolus subnetwork removed (C) from the gut microbiome of *Dendrobates imitator* and *Dendrobates monticola* species. The networks exhibit an organic structure, with edge weight represented by edge thickness, node degree by node size, and phyla color-coded.

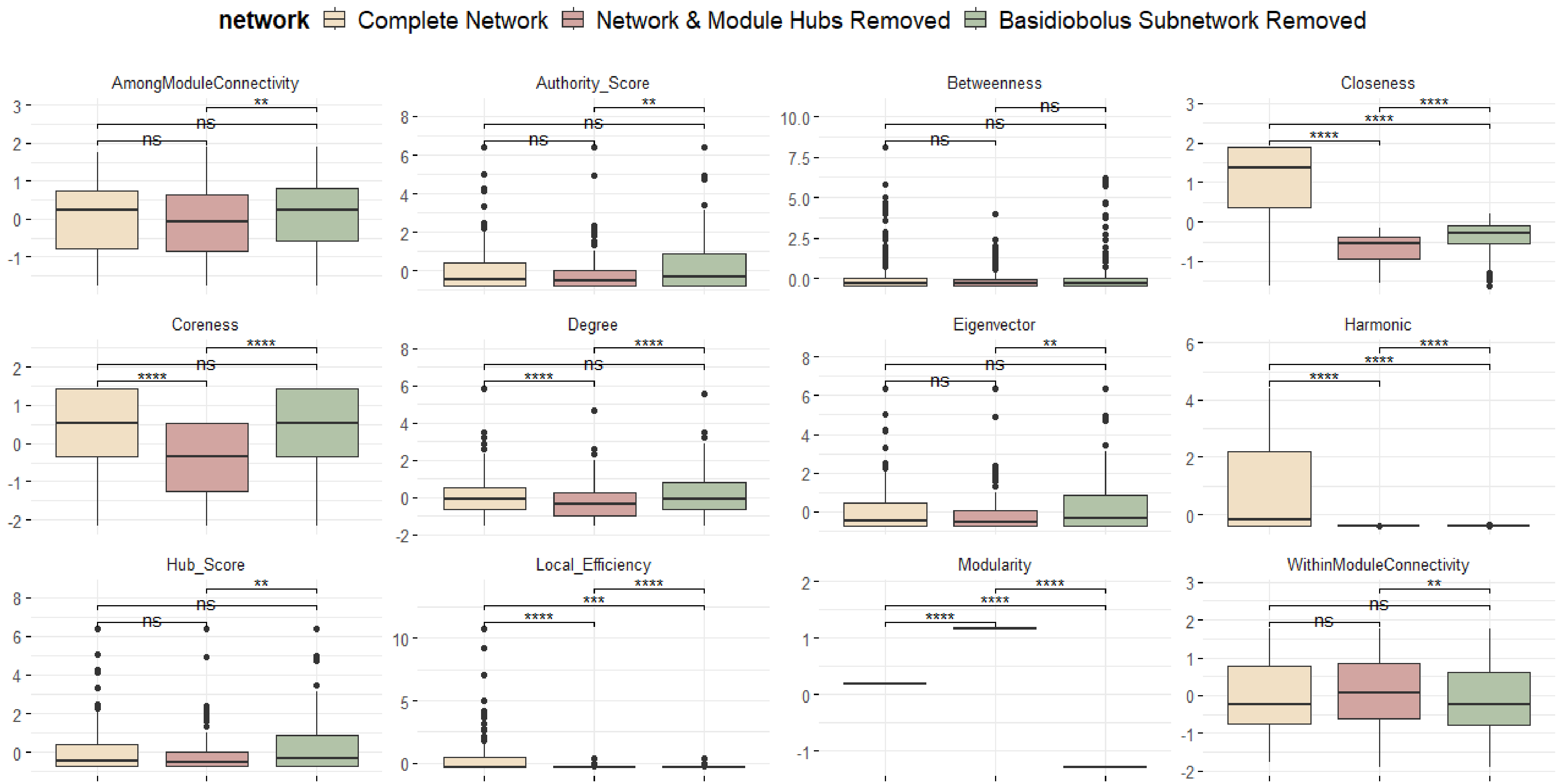


Figure S11 Comparison of network topographic properties across different conditions: (1) complete network, (2) network with module hubs removed, and (3) network with the *Basidiobolus* subnetwork removed.

### ITS scaling factor calculation

```
setwd("C:/Users/mghotbi/Documents/ForGitHub/Version1/ITSOTU")
physeq_ITSOTU <- readRDS("physeq_ITSOTU.rds")

library(DspikeIn)

## Thank you for using the DspikeIn package.

## For support, please contact Mitra Ghotbi at.

physeq_ITSOTU<-tidy_phyloseq(physeq_ITSOTU)

spiked_cells <- 733
species_name <- spiked_species<-merged_spiked_species<-"Dekkera_bruxellensis"
Dekkera <- subset_taxa(physeq_ITSOTU, Species=="Dekkera_bruxellensis")
hashcodes <- row.names(phyloseq::tax_table(Dekkera))

# Subset spiked samples (264 samples are spiked)
physeq_ITSOTU <- subset_samples(physeq_ITSOTU, spiked.volume %in% c("2", "1"))

Spiked_ITS_sum_scaled <- Pre_processing_species(
  physeq_ITSOTU,
  species_name,
  merge_method = "sum",
  output_file = "merged_physeq_sum.rds")

## Starting pre-processing...

## Processing species: Dekkera_bruxellensis

## ASVs found for species Dekkera_bruxellensis: 3f4a791b8dc05a20b40c164e663de4a0,
55e8f49352ccbdbf1c07f45d64d1ed75, 421e43995f1915381f1d96bfd32fe411, ed0d24e37e5032de032665488020762d,
90796b71f3e313827edd42e956a142a1

## Merged phyloseq object by summing abundances for species: Dekkera_bruxellensis

## Merged phyloseq object saved to: merged_physeq_sum.rds

## OTU table dimensions: 11862, 240

## Taxonomy table dimensions: 11862, 7

## OTU table row names: b0d8709d00c52cd6f95e80208ec58bed, 8802fa20235c7208c28a77bf31b16ab0,
7f78e32ee8be7c4e84c2468a78e56ed1, e6a1c236f2da2bb42b7863b40b151857, 4bb6692c6d9f4a818a80b78eefb05ee8,
4dccbfe4f811fe0d5871482a35c0256e

## Taxonomy table row names: b0d8709d00c52cd6f95e80208ec58bed, 8802fa20235c7208c28a77bf31b16ab0,
7f78e32ee8be7c4e84c2468a78e56ed1, e6a1c236f2da2bb42b7863b40b151857, 4bb6692c6d9f4a818a80b78eefb05ee8,
4dccbfe4f811fe0d5871482a35c0256e

## Pre-processing complete.

result <- calculate_spikeIn_factors(Spiked_ITS_sum_scaled, 733, merged_spiked_species)
```

```
## Identifying spiked species...
## Saved file: spikeIn_factors/Physeq_NO_Spiked_Species.rds
## Saved file: spikeIn_factors/Spiked_16S_Total_Reads.csv

## Warning in prune_taxa(taxa, phy_tree(x)): prune_taxa attempted to reduce tree to 1 or fewer tips.
##  tree replaced with NULL.

## Saved file: spikeIn_factors/Spiked_Species_MergedToOne.rds
## Saved file: spikeIn_factors/Spiked_Species_Reads.csv
## Negative values detected in spiked species reads. Converting to zero.
## Scaling factors saved as spikeIn_factors.csv.
## Process complete. Scaling factors for spiked species saved in the output directory.
## Table saved in docx format: spikeIn_factors/spikeIn_factors_summary.docx

scaling_factors<-result$scaling_factors

# Convert relative counts data to absolute counts
absolute <- convert_to_absolute_counts(Spiked_ITS_sum_scaled, scaling_factors)

## Absolute count data saved to:
C:/Users/mghotbi/Documents/ForGitHub/Version1/ITSOTU/physeq_adj_scaled_AbsoluteCount.csv

absolute_counts <- absolute$absolute_counts
physeq_absolute_abundance_ITS_OTU <- absolute$physeq_obj

# summary statistics
post_eval_summary <- calculate_summary_stats_table(absolute_counts)

##
## Attaching package: 'dplyr'

## The following objects are masked from 'package:stats':
##
##     filter, lag

## The following objects are masked from 'package:base':
##
##     intersect, setdiff, setequal, union

## Warning: package 'flextable' was built under R version 4.3.3

## Table saved in docx format: post_eval_summary.docx
## Summary statistics saved as CSV: post_eval_summary.csv

print(post_eval_summary)

## a flextable object.
## col_keys: `spiked.blank.20529_S180_mean`, `spiked.blank.20913_S180_mean`, …..

max_passed_range <- 11
output_path <- "spike_success_report.docx"
physeq_ITS_adj_scaled_perc <- subset_samples(physeq_absolute_abundance_ITS_OTU, sample.or.blank != "blank")
conclusion(physeq_ITS_adj_scaled_perc, merged_spiked_species, max_passed_range, output_path)

## Warning in prune_taxa(taxa, phy_tree(x)): prune_taxa attempted to reduce tree to 1 or fewer tips.
##  tree replaced with NULL.
```

```
## Table saved in docx format: spike_success_report.docx
## Merged data saved as CSV: spike_success_report.csv
## [1] "Structure of spike_success_report:"
## 'data.frame':    236 obs. of  5 variables:
##  $ Sample           : chr  "STP1719.20518_S143" "STP213.20519_S155" "STP268.20520_S167" "STP544.20515_S107"
...
##  $ Total_Reads_total : num  25451 24670 13414 28690 39910 ...
##  $ Total_Reads_spiked: num  7455 39 660 25971 13722 ...
##  $ Percentage       : num  29.292 0.158 4.92 90.523 34.382 ...
##  $ Result           : chr  "failed" "passed" "passed" "failed" ...
## NULL
## [1] "First few rows of spike_success_report:"
##              Sample Total_Reads_total Total_Reads_spiked Percentage Result
## 1 STP1719.20518_S143             25451               7455 29.2915799 failed
## 2  STP213.20519_S155             24670                 39  0.1580867 passed
## 3  STP268.20520_S167             13414                660  4.9202326 passed
## 4  STP544.20515_S107             28690              25971 90.5228303 failed
## 5  STP570.20516_S119             39910              13722 34.3823603 failed
## 6  STP579.20517_S131             99726              83429 83.6582235 failed
## [1] "Unique values in the 'Result' column after filtering and converting to lowercase:"
## [1] "failed" "passed"
## Table saved in docx format: spike_success_report.docx
## Summary statistics saved as CSV: spike_success_report.csv

##   mean_total_reads_spiked sd_total_reads_spiked se_total_reads_spiked
## 1                5942.077              20075.04              1312.346
##   q25_total_reads_spiked median_total_reads_spiked q75_total_reads_spiked
## 1                  147.5                       701                   3436
##   mean_percentage sd_percentage se_percentage q25_percentage median_percentage
## 1         19.3335      26.58252      1.737754       1.224706          6.244537
##   q75_percentage passed_count failed_count
## 1       26.97019          127          107
```

### scaling factor calculation for 16S rRNA

```
setwd("~/ForGitHub/Version1/16SOTU")
physeq_16SOTU <- readRDS("physeq_16SOTU.rds")

library(DspikeIn)

## Thank you for using the DspikeIn package.

## For support, please contact Mitra Ghotbi at.

spiked_cells <-1847
species_name <- spiked_species <- c("Tetragenococcus_halophilus", "Tetragenococcus_sp")
merged_spiked_species<-"Tetragenococcus_halophilus"
physeq_16SOTU<- DspikeIn::tidy_phyloseq(physeq_16SOTU)
Tetra <- phyloseq::subset_taxa(physeq_16SOTU,Species=="Tetragenococcus_halophilus" |  Species=="Tetragenococcus_sp")

## Warning in prune_taxa(taxa, phy_tree(x)): prune_taxa attempted to reduce tree to 1 or fewer tips.
##  tree replaced with NULL.
```

```
hashcodes <- row.names(phyloseq::tax_table(Tetra))

# Subset spiked samples (264 samples are spiked)
spiked_16S_OTU <- subset_samples(physeq_16SOTU, spiked.volume %in% c("2", "1"))

Spiked_16S_sum_scaled <- Pre_processing_species(
  spiked_16S_OTU,
  species_name,
  merge_method = "sum",
  output_file = "merged_physeq_sum.rds")

## Starting pre-processing...

## Processing species: Tetragenococcus_halophilus

## ASVs found for species Tetragenococcus_halophilus: 2ddb215ff668b6a24a9ccf2bb076a453

## No need to merge; only one ASV for species: Tetragenococcus_halophilus

## Processing species: Tetragenococcus_sp

## ASVs found for species Tetragenococcus_sp:

## No need to merge; only one ASV for species: Tetragenococcus_sp

## OTU table dimensions: 9334, 264

## Taxonomy table dimensions: 9334, 7

## OTU table row names: 020e00d90ba97c5898944ab6f7b1b7c9, b00466354053c9065c8aa3d6fbb33eaa,
f872c4bf84bcf44434fa2023788f6517, df13f71584d4a579c81d909eaba11a74, ed285eb1aac505a1f062b482300b69f7,
63f5509575600a9e7afb6847d6296976

## Taxonomy table row names: 020e00d90ba97c5898944ab6f7b1b7c9, b00466354053c9065c8aa3d6fbb33eaa,
f872c4bf84bcf44434fa2023788f6517, df13f71584d4a579c81d909eaba11a74, ed285eb1aac505a1f062b482300b69f7,
63f5509575600a9e7afb6847d6296976

## Pre-processing complete.

result <- calculate_spikeIn_factors(Spiked_16S_sum_scaled, 1847, merged_spiked_species)

## Identifying spiked species...
## Saved file: spikeIn_factors/Physeq_NO_Spiked_Species.rds
## Saved file: spikeIn_factors/Spiked_16S_Total_Reads.csv

## Warning in prune_taxa(taxa, phy_tree(x)): prune_taxa attempted to reduce tree to 1 or fewer tips.
##  tree replaced with NULL.

## Saved file: spikeIn_factors/Spiked_Species_MergedToOne.rds
## Saved file: spikeIn_factors/Spiked_Species_Reads.csv
## Negative values detected in spiked species reads. Converting to zero.
## Scaling factors saved as spikeIn_factors.csv.
## Process complete. Scaling factors for spiked species saved in the output directory.
## Table saved in docx format: spikeIn_factors/spikeIn_factors_summary.docx

scaling_factors<-result$scaling_factors

# Convert relative counts data to absolute counts
physeq_16S_adj_scaled_AbsoluteCount <- convert_to_absolute_counts(Spiked_16S_sum_scaled, scaling_factors)

## Absolute count data saved to: C:/Users/mghotbi/Documents/ForGitHub/Version1/16SOTU/physeq_adj_scaled_AbsoluteCount.csv

absolute <- convert_to_absolute_counts(Spiked_16S_sum_scaled, scaling_factors)
```

```
## Absolute count data saved to: C:/Users/mghotbi/Documents/ForGitHub/Version1/16SOTU/physeq_adj_scaled_AbsoluteCount.csv

absolute_counts <- physeq_16S_adj_scaled_AbsoluteCount$absolute_counts
physeq_absolute_abundance_16S_OTU <- physeq_16S_adj_scaled_AbsoluteCount$physeq_obj


physeq_absolute16S_OTU<-tidy_phyloseq(physeq_absolute_abundance_16S_OTU)

# summary statistics
post_eval_summary <- calculate_summary_stats_table(absolute_counts)

##
## Attaching package: 'dplyr'

## The following objects are masked from 'package:stats':
##
##     filter, lag

## The following objects are masked from 'package:base':
##
##     intersect, setdiff, setequal, union

## Warning: package 'flextable' was built under R version 4.3.3

## Table saved in docx format: post_eval_summary.docx
## Summary statistics saved as CSV: post_eval_summary.csv

print(post_eval_summary)

## a flextable object.
## col_keys: `spiked.blank.20433_S84_mean`, `spiked.blank.20817_S84_mean`, `Std2uL.20625_S84_mean`,
`StdSwab1uL.20624_S72_mean`, `STP1719.20422_S47_mean`, `STP213.20423_S59_mean`, `STP268.20424_S71_mean`,
`STP544.20419_S11_mean`,
```

```
> sessionInfo()

attached packages:
[1] DspikeIn_0.0.0.9000

loaded via a namespace (and not attached):
 [1] ade4_1.7-22          tidyselect_1.2.1     dplyr_1.1.4          Biostrings_2.70.1    bitops_1.0-7
 [6] fastmap_1.1.1        RCurl_1.98-1.13      phyloseq_1.46.0      promises_1.2.1       digest_0.6.33
[11] mime_0.12            lifecycle_1.0.4      cluster_2.1.6        ellipsis_0.3.2       survival_3.5-7
[16] magrittr_2.0.3       compiler_4.3.2       rlang_1.1.4          tools_4.3.2          igraph_1.6.0
[21] utf8_1.2.4           data.table_1.14.10   htmlwidgets_1.6.4    pkgbuild_1.4.4       plyr_1.8.9
[26] rsconnect_1.3.1      pkgload_1.4.0        Rtsne_0.17           miniUI_0.1.1.1       purrr_1.0.2.9000
[31] BiocGenerics_0.48.1  grid_4.3.2           stats4_4.3.2         fansi_1.0.6          urlchecker_1.0.1
[36] profvis_0.3.8        multtest_2.58.0      biomformat_1.30.0    xtable_1.8-4         colorspace_2.1-0
[41] Rhdf5lib_1.24.1      ggplot2_3.5.1        scales_1.3.0         iterators_1.0.14     MASS_7.3-60
[46] cli_3.6.1            vegan_2.6-4          crayon_1.5.3         remotes_2.5.0        generics_0.1.3
[51] microbiome_1.24.0    rstudioapi_0.16.0    reshape2_1.4.4       sessioninfo_1.2.2    ape_5.7-1
[56] cachem_1.0.8         rhdf5_2.46.1         stringr_1.5.1        zlibbioc_1.48.0      splines_4.3.2
[61] parallel_4.3.2       XVector_0.42.0       vctrs_0.6.5          devtools_2.4.5       Matrix_1.6-4
[66] jsonlite_1.8.8       IRanges_2.36.0       S4Vectors_0.40.2     foreach_1.5.2        tidyr_1.3.0
[71] glue_1.6.2           codetools_0.2-20     stringi_1.8.2        gtable_0.3.5         later_1.3.2
[76] GenomeInfoDb_1.38.7  munsell_0.5.1        tibble_3.2.1         pillar_1.9.0         htmltools_0.5.7
[81] rhdf5filters_1.14.1  GenomeInfoDbData_1.2.11 R6_2.5.1            shiny_1.8.1.1        lattice_0.22-5
[86] Biobase_2.62.0       memoise_2.0.1        httpuv_1.6.13        Rcpp_1.0.11          nlme_3.1-164
[91] permute_0.9-7        mgcv_1.9-0           usethis_2.2.3        fs_1.6.3             pkgconfig_2.0.3
```

To demonstrate the utility of the `DspikeIn` package;

```
# make sure about the presence of 'spiked.volume' column in the metadata
# 16s rRNA
spiked_cells <-1847
species_name <- spiked_species <- c("Tetragenococcus_halophilus", "Tetragenococcus_sp")
merged_spiked_species<-"Tetragenococcus_halophilus"
physeq_16SOTU<- DspikeIn::tidy_phyloseq(physeq_16SOTU)
Tetra <- phyloseq::subset_taxa(physeq_16SOTU,Species=="Tetragenococcus_halophilus" |
Species=="Tetragenococcus_sp")
hashcodes <- row.names(phyloseq::tax_table(Tetra))

# ITS rDNA
spiked_cells <- 733
species_name <- spiked_species<-merged_spiked_species<-"Dekkera_bruxellensis"
Dekkera <- subset_taxa(physeq_ITSOTU, Species=="Dekkera_bruxellensis")
hashcodes <- row.names(phyloseq::tax_table(Dekkera))
```

*Prerequisite list of species:*

```
# tidy up
physeq <- tidy_phyloseq(physeq)
spiked_species_list <- c("Pseudomonas aeruginosa", "Escherichia coli", "Clostridium difficile")
spiked_cells_list <- c(10000, 20000, 15000)
```

2.4.1 Preprocessing data

2.4.1 Preprocessing data

```
Single spiked species
 16S rRNA
 presence of 'spiked.volume' column in metadata
 spiked_cells <-1847
 species_name <- spiked_species <- c("Tetragenococcus_halophilus", "Tetragenococcus_sp")
 merged_spiked_species<-"Tetragenococcus_halophilus"
 Tetragenococcus <- phyloseq::subset_taxa(physeq_16SASV,
 Species=="Tetragenococcus_halophilus" | Species=="Tetragenococcus_sp")
 hashcodes <- row.names(phyloseq::tax_table(Tetragenococcus)
```

*List of spiked species*

```
spiked_species <- c("Pseudomonas aeruginosa", "Escherichia coli", "Clostridium difficile")
merged_physeq_sum <- Pre_processing_species_list(physeq, spiked_species, merge_method = "sum")
```

```
# tidy up
physeq <- tidy_phyloseq(physeq)
```

```
spiked_species_list <- c("Pseudomonas aeruginosa", "Escherichia coli", "Clostridium difficile")
spiked_cells_list <- c(10000, 20000, 15000)
merged_physeq_sum <- Pre_processing_species_list(physeq, spiked_species_list, merge_method = "sum")
```

*2.4.2 Scaling factors calculation*

*One species and equal spiked.in volume*

```
merged_spiked_species <- c("Tetragenococcus_halophilus")
# # Calculate scaling factors and generate the report
result <- calculate_spikeIn_factors(merged_physeq_sum, 1874, merged_spiked_species)
# # Access the results
scaling_factors <- result$scaling_factors
 physeq_no_spiked <- result$physeq_no_spiked
 spiked_16S_total_reads <- result$spiked_16S_total_reads
 spiked_species <- result$spiked_species
spiked_species_merged <- result$spiked_species_merged
spiked_species_reads <- result$spiked_species_reads
```

*List of species and variable spiked.in volume*

```
spiked_species_list <- list(
c("Pseudomonas aeruginosa"),
c("Escherichia coli"),
c("Clostridium difficile"))
spiked_cells_list <- c(10000, 20000, 15000)
scaling_factors <- calculate_list_average_scaling_factors(physeq, spiked_species_list,
spiked_cells_list, merge_method = "sum") print(scaling_factors)
```

```
calculate_spike_percentage_list(merged_physeq_sum, merged_spiked_species = spiked_species_list, passed_range =
c(0.1, 20))
```

*2.4.3 Converting to absolute abundance*

```
absolute <- convert_to_absolute_counts(merged_physeq_sum, scaling_factors)
absolute_counts <- absolute$absolute_counts
physeq_obj <- absolute$physeq_o
```

*2.4.4 Normalization*

```
 ps= physeq_absolute_abundance_16S_ perc
# Example for TSS normalization
ps<-remove_zero_negative_count_samples(ps)
Host.species <- as.factor(ps@sam_data$Host.species)
result_tss <- normalization_set(ps, method = "tss")
```

```
# Extract the normalized dataset from the result
normalized_ps_tss <- result_tss$dat.normed
# Retrieve the scaling factors used during the TSS normalization
scaling_factors_tss <- result_tss$scaling.factor
# 'normalized_ps_tss' now contains the normalized data,
# and 'scaling_factors_tss' holds the scaling factors applied during normalization.

# Example for Poisson normalization
result_Poisson <- normalization_set(ps,
method = "Poisson", groups = "Host.genus")
normalized_ps_Poisson <- result_Poisson$dat.normed
scaling_factors_Poisson <- result_Poisson$scaling.factor


# Example for TMM normalization
result_TMM <- normalization_set(ps,
method = "TMM", groups = "Animal.type")
normalized_ps_TMM <- result_TMM$dat.normed
scaling_factors_TMM <- result_TMM$scaling.factor
# Example for TC normalization

result_TC <- normalization_set(ps, method = "TC", groups = "Host.species")
normalized_ps_TC <- result_TC$dat.normed
scaling_factors_TC <- result_TC$scaling.factor

# Example for Quantile normalization
result_QN <- normalization_set(ps, method = "QN")
normalized_ps_QN <- result_QN$dat.normed
scaling_factors_QN <- result_QN$scaling.factor

# Example for UQ normalization
result_UQ <- normalization_set(ps, method = "UQ", groups = "Host.species")
normalized_ps_UQ <- result_UQ$dat.normed
scaling_factors_UQ <- result_UQ$scaling.factor


# Example for Median normalization
result_med <- normalization_set(ps,
method = "med", groups = "Host.species")
normalized_ps_med <- result_med$dat.normed
scaling_factors_med <- result_med$scaling.factor

# Example for CLR normalization
result_clr <- normalization_set(ps, method = "clr")
normalized_ps_clr <- result_clr$dat.normed
scaling_factors_clr <- result_clr$scaling.factor

# Example for Rarefying
result_rar <- normalization_set(ps, method = "rar")

normalized_ps_rar <- result_rar$dat.normed
scaling_factors_rar <- result_rar$scaling.factor
```

*2.4.5 Differential abundance and Random Forest*

```
results_edgeR <- perform_and_visualize_DA_edgeR(
  ps = abs_monticola_imitator,
  group_var = "Host.taxon",
  contrast = c("Desmognathus monticola", "Desmognathus imitator"),
  output_csv_path = "DA_edgeR.csv",
  target_glom = "Genus",
  significance_level = 0.05)

## Removing 7074 features with zero counts across all samples.
## Column names in the design matrix:
##  Desmognathus.imitator Desmognathus.monticola
## edgeR results saved to: DA_edgeR.csv

# Visualize Volcano Plot
print(results_edgeR$plot)
results_edgeR$results
results_edgeR$ps_significant

rf_physeq <- RandomForest_selected_ASVs group_var ( absolute ,response_var = " Host.taxon ",
na_vars = c("Habitat", " Ecoregion.III", " Host.taxon ", " Diet"))
saveRDS(rf_physeq,"rf_physeq.rds")
```

*2.4.6 Visualization*

```
# Plot a glommed tree with specified resolution
physeq_ITSOTU <- readRDS("physeq_ITSOTU.rds")
Dekkera <- phyloseq::subset_taxa(physeq_ITSOTU, Species=="Dekkera_bruxellensis")
DspikeIn::plot_glommed_tree(Dekkera, resolution = 0.0)

# Custom tree plotting with circular layout
DspikeIn::plot_tree_custom(Dekkera, output_prefix = "p0", width = 18, height = 18, layout = "circular")

# Bootstrap tree with cophenetic distances and save as PNG
DspikeIn::Bootstrap_phy_tree_with_cophenetic(physeq_object = Dekkera, output_file =
"tree_with_bootstrap_and_cophenetic.png", bootstrap_replicates = 1000)
# Define path to the FASTA file
fasta_path <- "~/DspikeIn_R /tetra.fasta"
# Plot neighbor-joining tree and save to file
DspikeIn::plot_tree_nj(fasta_path, output_file = "neighbor_joining_tree_with_bootstrap.png")
# Plot tree with alignment and save with specified dimensions
DspikeIn::plot_tree_with_alignment(Dekkera, output_prefix = "tree_alignment", width = 15, height = 15)

absolute=physeq_absolute_abundance_16S_ perc
abs_BlueRidge<-DspikeIn::ridge_plot_it(absolute, taxrank = "Genus",top_n = 20)
+ scale_fill_manual(values = color_palette$light_MG)

# Melt the phyloseq object to a long-format data frame
 pps_Abs<-psmelt(absolute)
```

```
alluvial_plot_abs <- alluvial_plot(  data = pps_Abs,
  axes = c( "Clade.Order","Animal.ecomode","Ecoregion.III","Diet","Reproduction","Metamorphosis"),
  abundance_threshold = 3000,
   fill_variable = "Genus",
  silent = TRUE,
  abundance_type = "absolute",
  top_taxa = 20,
  text_size = 4,
  legend_ncol = 1,
  custom_colors =   color_palette$light_MG)

# Define custom detection thresholds
custom_detections <- 10^seq(log10(3e-1), log10(0.5), length = 5)
# Plot core microbiome at the "Family" level
 PCM <- plot_core_microbiome_custom(
  absolute,
  detections = custom_detections,
  taxrank = "Family",
  output_core_rds = "core_microbiome.rds",
  output_core_csv = "core_microbiome.csv")

# Generate a barplot for absolute abundance at the genus level
 bp_ab <- taxa_barplot(
 physeq = absolute,  # phyloseq object
 target_glom = "Genus",
 treatment_variable = "Phylum",
 abundance_type = "absolute",
 x_angle = 0,
 fill_variable = "Genus",
 facet_variable = "Host.species")

# Adding custom theme and color scale
bp_ab$barplot +
  my_custom_theme() +
  xlab(NULL) +
  theme(axis.text.x = element_text(angle = 90, hjust = 1)) +
 scale_fill_manual(values = color_palette$light_MG)

taxa_barplot(
 physeq = absolute,  # phyloseq object
 target_glom = "Genus",
 treatment_variable = "Phylum",
 abundance_type = "absolute",

regression_plot(
data = metadata,
x_var =  " Richness",   # metadata is a data frame format
y_var =  "Total_Reads_spiked",
 custom_range =  c(0.1, 15, 30, 50, 80, 100),  # Define percentage ranges
 plot_title = NULL )
print(plot_object)
```
